# ACLY integrates metabolism and chromatin accessibility to enable B Cell activation and humoral immunity

**DOI:** 10.64898/2026.05.24.727510

**Authors:** Meilu Li, Zhiqiang Zhang, Xian Zhou, Yanfeng Li, Xingxing Zhu, Aditya Bhagwate, Nagaswaroop Kengunte Nagaraj, Hu Zeng

## Abstract

B cell activation and differentiation into antibody-secreting cells require extensive metabolic and epigenetic remodeling, yet the molecular mechanisms that integrate these programs remain incompletely understood. ATP-citrate lyase (ACLY) links glucose metabolism to acetyl-CoA production, supporting lipid biosynthesis and protein acetylation. However, its role in humoral immunity has not been fully defined. Here, using genetic and integrated multi-omics approaches, we show that B cell activation is accompanied by coordinated metabolic, transcriptional, and epigenetic reprograming. Although ACLY is dispensable for B cell development and homeostasis, it is required to establish chromatin accessibility programs in activated B cells, with a more pronounced impact on the epigenetic landscape than on transcriptional output. ACLY-deficient B cells exhibit profound defects in TLR and BCR elicited activation, survival and metabolic fitness *ex vivo*. *In vivo*, B cell-intrinsic loss of ACLY results in impaired antigen-specific antibody production, associated with reduced germinal center and plasmablast formation, but normal homeostatic proliferation. Deletion of ACLY after B cell activation reduces plasmablast generation *in vivo*, indicating a continued requirement for ACLY beyond the initial activation phase. Together, these findings identify ACLY as a central regulator that links metabolism to epigenetic programing that supports B cell activation and humoral immunity.

## Introduction

Lipid biosynthesis plays a pivotal role in cellular metabolism and immunity^1^. The direct precursor of lipid biosynthesis is acetyl-coenzyme A (acetyl-CoA), which is also essential for bioenergetics, nucleotide biosynthesis, protein acetylation and epigenetic regulation. Production of acetyl-CoA is compartmentalized. Mitochondrial acetyl-CoA is produced by pyruvate dehydrogenase and is condensed with oxaloacetate by citrate synthase to form citrate, which enters tricarboxylic acid (TCA) cycle. Alternatively, citrate can be exported to cytosol where it is split into oxaloacetate and acetyl-CoA by ATP citrate lyase (ACLY)^2^. Cytosolic acetyl-CoA can also be produced from acetate by Acetyl-CoA synthetase 2 (ACSS2)^3^. ACLY controls histone acetylation and lipogenesis in a variety of mammalian cells^4–6^. Within immune cells, ACLY promotes M2 activation of macrophages^7^, and memory CD8 T cell function^8^. ACLY and ACSS2 cooperatively promote CD8 T cell effector responses^9^, but have divergent roles in modulating CD8 T cell exhaustion and anti-tumor immunity^10^. Furthermore, ACLY is considered dispensable for early T cell activation, because upon anti-CD3/anti-CD28 stimulation, ACLY-deficient T cells had normal expression of Ki67, multiple activation markers, and only a modest reduction in cell division^11^. Previous study indicates that LPS stimulation increases ACLY activity in B cells and pharmacological inhibition of ACLY reduces B cell proliferation *in vitro* associated with reduced fatty acid synthesis^12^. Yet, the functions of ACLY in B cell development, homeostasis, activation, and humoral immunity remain poorly defined.

Herein, we use genetic models in which *Acly* is deleted in lymphocytes or activated B cells to examine their contributions to B cell biology. Our data demonstrate that ACLY is essential for B cell activation, survival and metabolism *ex vivo*, yet it is not required for development or homeostasis at steady state. B cell-intrinsic ACLY is required for optimal GC and plasmablast formation and for antibody production. Integrative multi-omics analyses reveal that B cell activation invokes coordinated metabolic, transcriptional and epigenetic reprograming. Surprisingly, ACLY deficiency has a disproportionate impact on the epigenetic landscape, leading to a global attenuation of chromatin accessibility with comparatively modest effects on transcription. Finally, we show that ACLY plays an important role beyond early B cell activation phase to sustain humoral immune responses. Collectively, our study establishes ACLY as a critical metabolic regulator that links acetyl-CoA production to chromatin remodeling, thereby supporting B cell activation and humoral immunity.

## Results

### Lymphocyte development and homeostasis are largely preserved after ACLY deletion

To establish the physiological context for ACLY function in B cells, we generated *Cd2*^iCre^*Acly*^fl/fl^ mice, in which *Acly* was deleted specifically in lymphocytes. Loss of *Acly* mRNA expression in B cells was confirmed by qRT-PCR (Supplementary Fig. 1A). Deletion of *Acly* did not perturb B cell development or steady-state homeostasis in the bone marrow (BM), spleen and peritoneal cavity (Supplementary Fig. 1B-D). Steady-state germinal center (GC) formation in mucosal-associated lymphoid tissues was preserved, as Peyer’s patches from *Cd2*^iCre^*Acly*^fl/fl^ mice displayed normal frequencies and numbers of GL7^+^Fas^+^ GC B cells (Supplementary Fig. 1E). Likewise, IgA^+^ B cells and CD138^+^TACI^+^ plasma cells in the colonic lamina propria were comparable between genotypes (Supplementary Fig. 1F and G). In contrast, BM CD138^+^TACI^+^ plasma cells showed a modest but reproducible reduction in *Cd2*^iCre^*Acly*^fl/fl^ mice (Supplementary Fig. 1H), suggesting a selective requirement for ACLY in sustaining BM plasma cell compartments. Despite the modest reduction of BM plasma cells, baseline serum immunoglobulin levels, including IgG subclasses, IgA, and IgM, were indistinguishable between WT and *Cd2*^iCre^*Acly*^fl/fl^ mice (Supplementary Fig. 1I).

In addition to B cell compartments, T cell development was largely preserved in *Cd2*^iCre^*Acly*^fl/fl^ mice, albeit with a stage-specific alteration in the thymus. Flow cytometry revealed comparable frequencies and absolute numbers of double-negative (DN) thymocytes between WT and *Cd2*^iCre^*Acly*^fl/fl^ mice, whereas double-positive (DP) thymocytes were modestly reduced in the absence of ACLY (Supplementary Fig. 1J). Despite this reduction, both CD4^+^ single-positive (CD4^+^ SP) and CD8^+^ single-positive (CD8^+^ SP) thymocyte populations were present at normal frequencies and cell numbers (Supplementary Fig. 1J). *Cd2*^iCre^*Acly*^fl/fl^ mice also had completely normal CD4^+^ T cells, including Foxp3^+^ regulatory T cells, and CD8^+^ T cells in the spleen and peripheral lymph nodes (Supplementary Fig. 1K and L), indicating that ACLY is dispensable for the generation and homeostasis of mature T cells. Together, these data demonstrate that although ACLY deficiency does not impair lymphocyte development, peripheral B and T cell homeostasis, or steady-state immunoglobulin levels.

### ACLY is selectively required for TLR and BCR engagement elicited B cell activation, proliferation, and survival but is dispensable for homeostatic proliferation *in vivo*

To determine whether ACLY directly supports B cell activation and effector responses, we first examined the regulation of ACLY and the alternative acetyl-CoA–producing enzyme ACSS2 during lymphocyte activation. Immunoblot analysis revealed a robust induction of ACLY following activation of splenic B cells with LPS/IL-4/BAFF, whereas changes in ACLY expression in activated CD4⁺ T cells were comparatively modest (Fig. 1A). In contrast, ACSS2 expression exhibited only subtle changes upon activation in both B cells and CD4⁺ T cells, suggesting that ACLY might be the predominant acetyl-CoA–producing enzyme dynamically regulated during B cell activation. Efficient deletion of ACLY protein in *Cd2*^iCre^*Acly*^fl/fl^ B cells was confirmed by immunoblot (Fig. 1B). ACLY-deficient B cells had a modest reduction of cellular acetyl-CoA level (Supplementary Fig. 2A), suggesting that loss of ACLY might not drastically change the total acetyl-CoA pool, consistent with previous report^9^. Upon stimulation, *Cd2*^iCre^*Acly*^fl/fl^ B cells exhibited significantly reduced upregulation of the activation marker CD69 and nutrient transporters CD71 and CD98 (Fig. 1C). We next tested whether alternative fuel source, acetate, and ACLY downstream product, oleic acid (OA), could rectify the activation defects of ACLY-deficient B cells. Supplementation with sodium acetate robustly restored the expression of all three activation markers, whereas OA provided only partial rescue (Fig. 1C). These results indicate that ACLY-dependent acetyl-CoA production is required for optimal B cell activation and that acetate can partially bypass this requirement.

**Figure 1.**
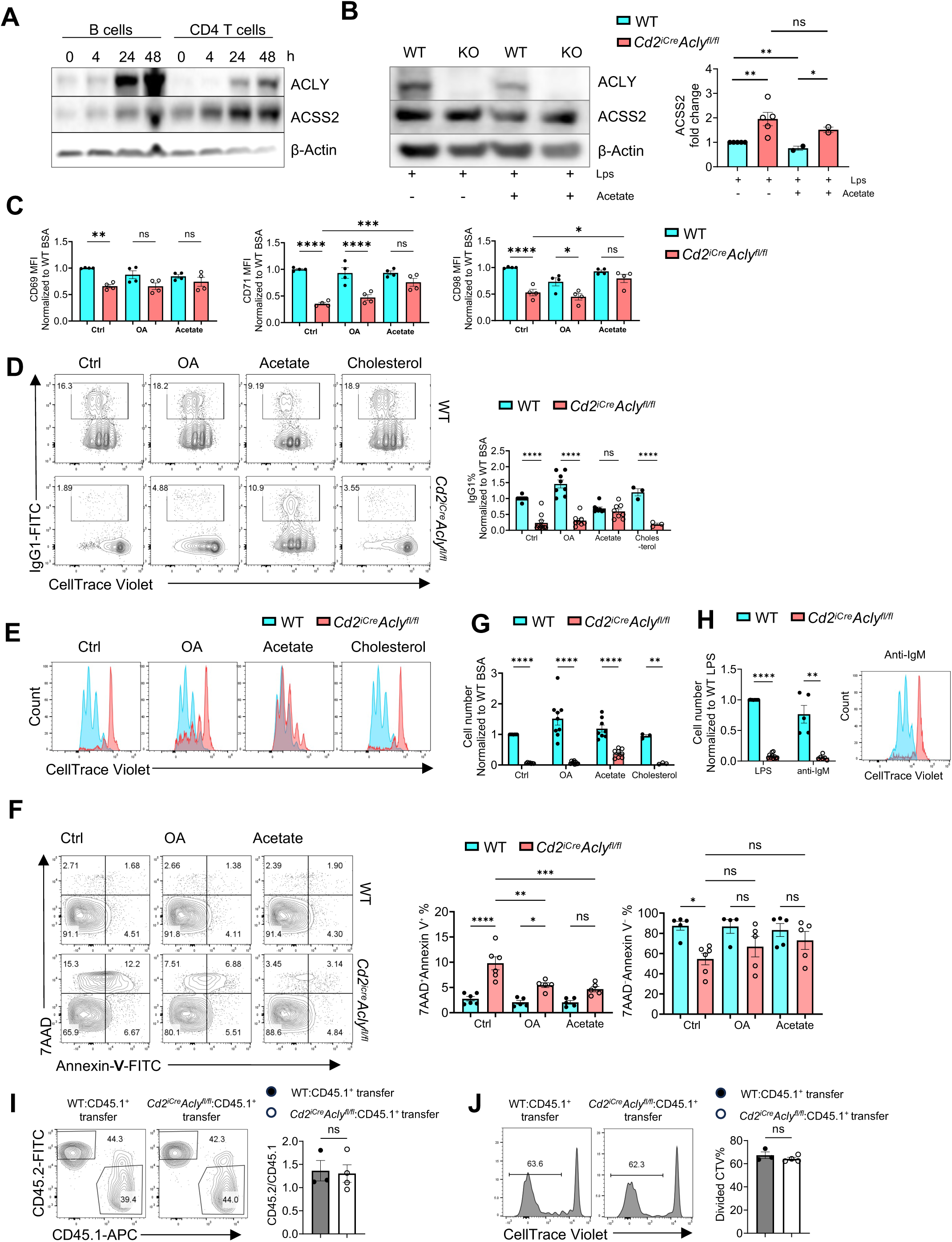
ACLY is critically required for B cell activation, proliferation and survival, but not homeostatic proliferation. (A) B cells and CD4^+^ T cells isolated from the spleens of B6 mice were activated with LPS/IL4/BAFF and anti-CD3/anti-CD28, respectively, at different time points. Immunoblot analysis of ACLY and ACSS2 were performed on B and CD4^+^ T cells, with β-Actin serving as the control. (B) Left, B cells from WT and *Cd2^iCre^Acly*^fl/fl^ mice stimulated LPS/IL4/BAFF, treated with or without sodium acetate for 16 h. Immunoblot analysis of ACLY, ACSS2. Right, quantification of the ACSS2 expression relative to that in WT B cells at baseline. (C) Normalized CD69, CD71 and CD98 mean fluorescence intensity (MFI) on LPS/IL4/BAFF activated B cell in the presence of BSA (50 µM) as control, OA (50 µM), or sodium acetate (5 mM) overnight. (D) B cells from WT and *Cd2^iCre^Acly*^fl/fl^ mice were labeled with Celltrace violet (CTV) and activated with LPS/IL4/BAFF in the presence of BSA, OA, sodium acetate, or cholesterol (5 μg/mL) for 3 days. Left, flow cytometry of CTV and IgG1 expression. Right, summary of IgG1^+^ percentages normalized to WT BSA group. (E) Cell proliferation measured by dilution of CTV LPS/IL4/BAFF activated B cells. (F) Left, flow cytometry of 7AAD and Annexin-V expression on LPS/IL4/BAFF activated B cells in the presence of BSA, OA, or sodium acetate overnight. Middle, the frequencies of dead cells (7AAD^+^Annexin-V^+^). Right, the frequencies of live cells (7AAD^−^Annexin-V^−^). (G) Cell numbers of B cells activated with LPS/IL4/BAFF in the presence of BSA, OA, sodium acetate, or cholesterol, for 3 days. Data was normalized to the number from BSA control condition, which was set as “1”. (H) Left, cell number of B cells activated with LPS/IL4/BAFF, or anti-IgM/IL4/BAFF for 3 days (normalized to the WT cells). Right, representative flow cytometry of CTV dilution in B cells activated with anti-IgM/IL4/BAFF. (I, J) B cells were purified from WT and *Cd2^icre^Acly^fl/fl^,* labled with CTV, mixed with B cells from CD45.1 congenic mice at 1:1 ratio, and transferred into *Rag1*^−/–^ mice. Left, expression of CD45.2 and CD45.1 in spleens of the recipicent mice at day 12 post transfer. Right, summary of the CD45.2/CD45.1 ratio (I). Flow cytometry of CTV expression in WT or ACLY-deficient donor cells. Right, summary of the frequencies of CTV^−^ populations. *p* values were calculated using Student’s t test. ns, not significant, *p < 0.05, **p < 0.01, ***p < 0.001, \*\*\*\**p* < 0.0001. Data are representative of at least three independent experiments (B-H). Error bars represent SEM.

To evaluate downstream functional consequences, we examined B cell proliferation and class-switch recombination. ACLY-deficient B cells displayed markedly reduced IgG1 expression and cell division compared with WT cells (Fig. 1D and E). Notably, sodium acetate restored IgG1 expression in ACLY-deficient B cells to levels comparable to the WT cells, whereas OA and cholesterol failed to rescue class-switch recombination. We next assessed cell survival. Annexin-V and 7AAD staining revealed increased frequencies of dead (7AAD⁺Annexin-V⁺) cells and decreased frequencies of live (7AAD⁻Annexin-V⁻) cells in the absence of ACLY, which were partially restored by either oleic acid or sodium acetate (Fig. 1F). Consistent with impaired proliferation and/or survival, total recovered B cell numbers after three days of culture were profoundly reduced in the absence of ACLY (Fig. 1G). Supplementation with sodium acetate, OA, or cholesterol failed to recover cell numbers (Fig. 1G). Reduced cell numbers were observed following both LPS/IL-4/BAFF and anti-IgM/IL-4/BAFF stimulation (Fig. 1H), indicating that ACLY supports B cell expansion downstream of both TLR4 and BCR pathways. Finally, to assess whether ACLY was required for homeostatic B cell proliferation *in vivo*, we transferred CellTrace Violet (CTV)-labeled WT or ACLY-deficient B cells, mixed with congenic CD45.1^+^ B cells at 1.1 ratio, into *Rag1*^−/–^ mice. We examined transferred donor cell CD45.2/CD45.1 ratio and cell division at 12 days post transfer. The ratio of CD45.2⁺ donor B cells to CD45.1⁺ competitor cells and cell division were both comparable between WT and ACLY-deficient B cells (Fig. 1I and J). Altogether, these data indicate that ACLY is selectively required for TLR/BCR-driven B cell activation, proliferation, and survival, but is dispensable for homeostatic proliferation in a lymphopenia environment.

To determine whether the impaired B cell activation following ACLY deletion resulted from defects in proximal B cell receptor (BCR) signaling, we examined early signaling events emanated from BCR ligation. Flow cytometric analysis of surface IgM internalization revealed no significant differences in kinetics or normalized IgM⁺ frequencies between WT and ACLY-deficient B cells at 0, 2, and 4 hours following anti-IgM stimulation (Supplementary Fig. 2B), indicating intact BCR engagement and internalization. Consistently, immunoblot analysis demonstrated comparable phosphorylation of key proximal BCR signaling components, including AKT (Ser473), BLNK, and SYK between WT and ACLY-deficient B cells following anti-IgM stimulation (Supplementary Fig. 2C). These data indicate that ACLY deficiency does not disrupt early BCR signaling cascades.

To assess whether ACLY deletion broadly affects T cell activation, CD4⁺ T cells from WT and *Cd2*^iCre^*Acly*^fl/fl^ mice were labeled with CTV and cultured in the presence of BSA, OA or sodium acetate. Under control (BSA-treated) conditions, ACLY-deficient CD4⁺ T cells exhibited modest but significant reductions in both proliferative frequency and total cell recovery, but no increased cell death, compared with WT controls. OA treatment partially normalized proliferative frequency but did not fully restore cell recovery, whereas sodium acetate treatment largely restored cell division and cell number recovery (Supplementary Fig. 2D). These findings are consistent with previous publication^11^, and contrast the exquisite dependence of ACLY for B cell activation and survival, suggesting a differential requirement of ACLY for early activation between B cells and T cells.

### ACLY supports metabolic reprogramming in activated B cells

Because ACLY is a key metabolic enzyme to produce cytosolic acetyl-CoA which feeds into tricarboxylic acid (TCA) cycle, we next investigated metabolic phenotypes of ACLY-deficient B cells. Extracellular flux analysis revealed that activated ACLY-deficient B cells exhibited significantly reduced extracellular acidification rate (ECAR) (Fig. 2A) and oxygen consumption rate (OCR) (Fig. 2B), indicating severely compromised glycolysis and mitochondrial oxidative phosphorylation. Treatment with sodium acetate partially rescued ECAR or OCR in ACLY-deficient B cells, whereas OA had a more modest effect (Fig. 2A,B), suggesting that acetate can support glycolytic flux and mitochondrial metabolism in the absence of ACLY but does not fully restore cellular metabolism.

**Figure 2.**
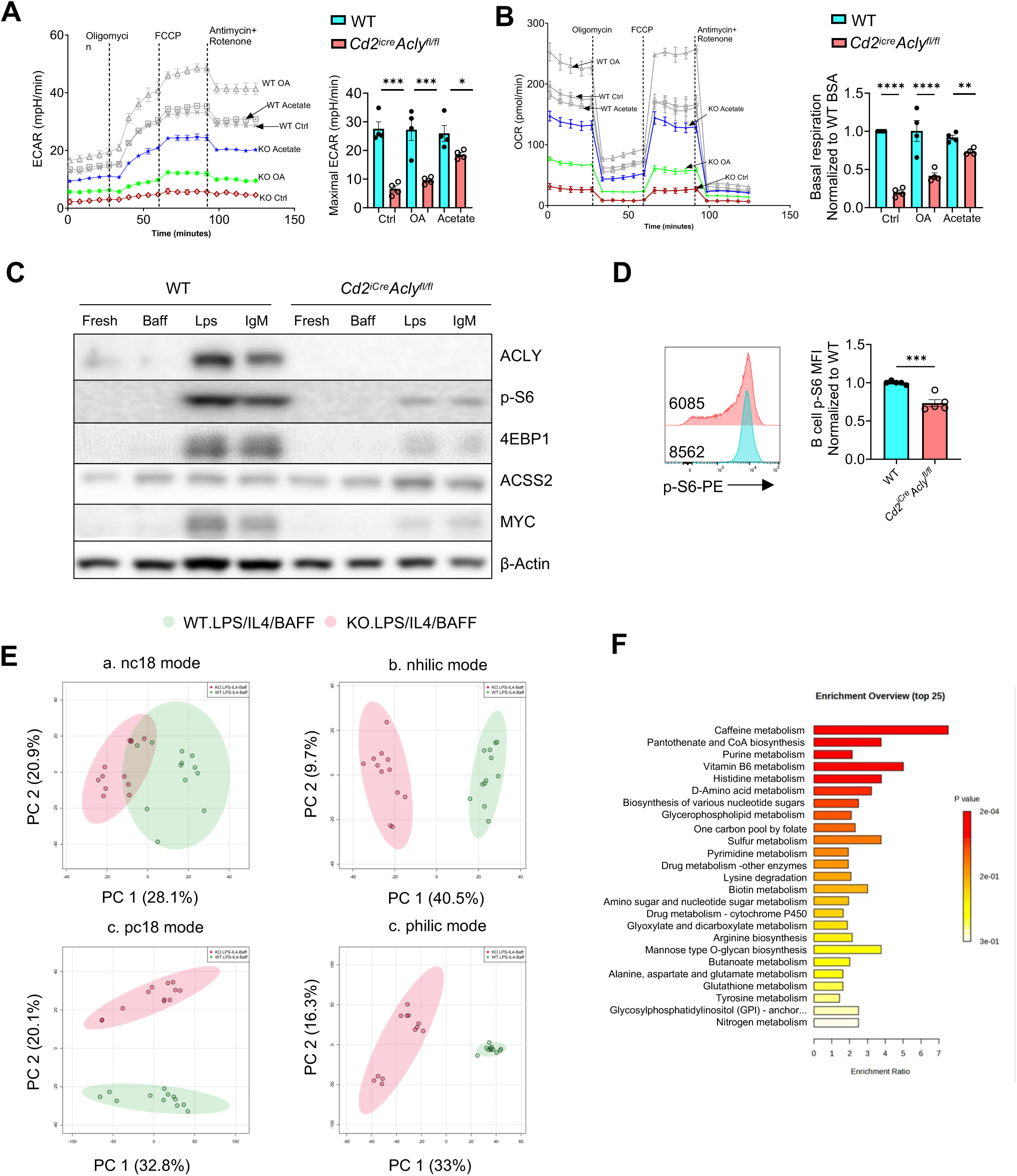
ACLY sustains metabolic reprograming in B cells. (A, B) B cells were activated with LPS/IL4/BAFF in the presence of BSA (50 µM), OA (50 µM), or sodium acetate (5mM). Equal number of live WT or ACLY-deficient B cells were used to measure extracellular acidification rate (ECAR) (A) and oxygen consumption rate (OCR) (B) on a Seahorse XF96 instrument. (A), Left, representative ECAR plot. Right: summary of maximal ECAR. (B) Left, representative OCR plot. Right, summary of basal respiration (normalized to WT B cells under control condition). (C) B cells from WT and *Cd2^iCre^Acly*^fl/fl^ mice were stimulated with BAFF, LPS/IL4/BAFF, or anti-IgM/IL4/BAFF overnight. Immunoblot analysis of ACLY, p-S6, 4EBP1, ACSS2 and total MYC were performed on these B cells, with β-Actin serving as the control. (D) Left, flow cytometry of p-S6 expression on activated B cell. Right, normalized p-S6 MFI. (E) Fresh and LPS/IL4/BAFF activated B cells were undergone untargeted metabolomics analysis. Principal component analysis (PCA) of metabolite profiles from stimulated WT and KO B cells. Metabolites were analyzed using four LC-MS acquisition modes: hydrophobic, “C18” and hydrophilic, “hilic”, each with positive (pc18, philic) and negative (nc18, nhilic) models. (F) Enrichment analysis on stimulated ACLY-deficient B cells vs WT B cells was performed using MetaboAnalyst. Data are representative of at least three independent experiments. p values were calculated using Student’s t-test. ns, not significant, *p < 0.05, **p < 0.01, ***p < 0.001, ****p < 0.0001. Error bars represent SEM.

B cell metabolic programing is controlled by the metabolic master regulator mTORC1 and transcription factor Myc^13–15^. Immunoblot demonstrated that under both LPS and anti-IgM activation, ACLY-deficient B cells exhibited reduced upregulation of the mTORC1 targets, p-S6, p-4EBP1, and MYC compared with WT cells (Fig. 2C). Flow cytometry further confirmed a marked reduction in p-S6 levels in activated ACLY-deficient B cells (Fig. 2D). These results indicate that ACLY is required for full activation of the mTORC1–MYC axis downstream of both TLR4 and BCR-dependent signals. In contrast to the pronounced signaling defects observed in B cells, mTORC1 activation in CD4^+^ and CD8^+^ T cells was relatively preserved (Supplementary Fig. 2E and F). Together, these data demonstrate that ACLY is required for coordinated glycolytic and mitochondrial metabolic reprogramming and for sustained mTORC1–MYC signaling in activated B cells.

Consistent with these metabolic defects, high-performance liquid chromatography (HPLC)-based untargeted metabolomic profiling revealed distinct metabolite alterations between WT and ACLY-deficient B cells (Fig. 2E and Supplementary Fig. 3A and B). Among the top discriminative features ranked by variable importance in projection (VIP), several metabolites associated with polyamine metabolism, amino acid metabolism and acyl-carnitine metabolism, including N1, N12-diacetylspermine, lysine, and carnitine, were prominently enriched (Supplementary Fig. 3C), suggesting perturbations in cellular acetylation potential and energy utilization. Pathway enrichment analysis further demonstrated that differentially abundant metabolites were significantly associated with multiple metabolic programs, including pantothenate and CoA biosynthesis, purine metabolism, D-amino acid metabolism, one-carbon metabolism via folate, arginine biosynthesis, and amino sugar and nucleotide sugar metabolism (Fig. 2F). These pathways collectively suggest a broad remodeling of central carbon metabolism and biosynthetic capacity upon ACLY deletion.

### ACLY couples acetyl-CoA metabolism to transcriptional and histone-acetylation programs upon B cell activation

In addition to supporting lipid metabolism, acetyl-CoA is also known to be used for epigenetic modifications and influences transcription^9,10^. To comprehensively assess ACLY-dependent transcriptional and epigenetic landscape, we performed RNA-seq and ATAC-seq on fresh and activated B cells from WT and *Cd2*^iCre^*Acly*^fl/fl^ mice (Fig. 3A). Principal component analysis (PCA) of bulk RNA-seq data revealed minimal transcriptional differences between WT and *Cd2*^iCre^*Acly*^fl/fl^ B cells under resting conditions (KO_Fresh vs WT_Fresh) (Fig. 3B). In contrast, upon stimulation (KO_Stim vs WT_Stim), WT and ACLY-deficient B cells separated distinctly, indicating substantial transcriptional reprogramming. Venn diagram comparison and volcano plots showed a markedly increased number of differentially expressed genes (DEGs) after stimulation compared to the resting state (Fig. 3C). However, the magnitude of stimulation induced transcriptional changes (both upregulation and downregulation, comparing stimulated to fresh) was largely comparable between WT and ACLY-deficient B cells (Fig. 3D). Functional enrichment analyses showed that activation of WT B cell induced cell cycle and metabolic programs, including chromosome segregation, mitotic cell cycle phase transition, as well as biosynthesis of cofactors and amino acids (Fig. 3E and F, Supplementary Fig. 5A and B). Conversely, genes reduced in KO_Stim were significantly enriched in pathways governing DNA replication and mitotic cell cycle phase transition (Fig. 3G; related analyses in Supplementary Fig. 5C and D). KEGG pathway analysis further demonstrated suppression of cell cycle and central metabolic programs, including carbon metabolism, nucleotide biosynthesis, purine and pyrimidine metabolism, and glycolysis/gluconeogenesis (Fig. 3H). Gene set enrichment analysis (GSEA) further revealed a shift in transcriptional programs, with ACLY-deficient cells exhibiting increased enrichment of inflammatory pathways, including interferon responses, p53 signaling, and apoptosis-related programs, alongside marked downregulation of proliferation/metabolism-associated pathways such as MYC targets, E2F targets and mTORC1 signaling (Fig. 3I). These transcriptional changes closely aligned with the profound proliferative and metabolic defects following ACLY deletion, indicating a level of coordinated disruption of central metabolic and biosynthetic programs. These findings indicate that ACLY is required to sustain proliferative and biosynthetic programs while restraining inflammatory and interferon responses during B cell activation.

**Figure 3.**
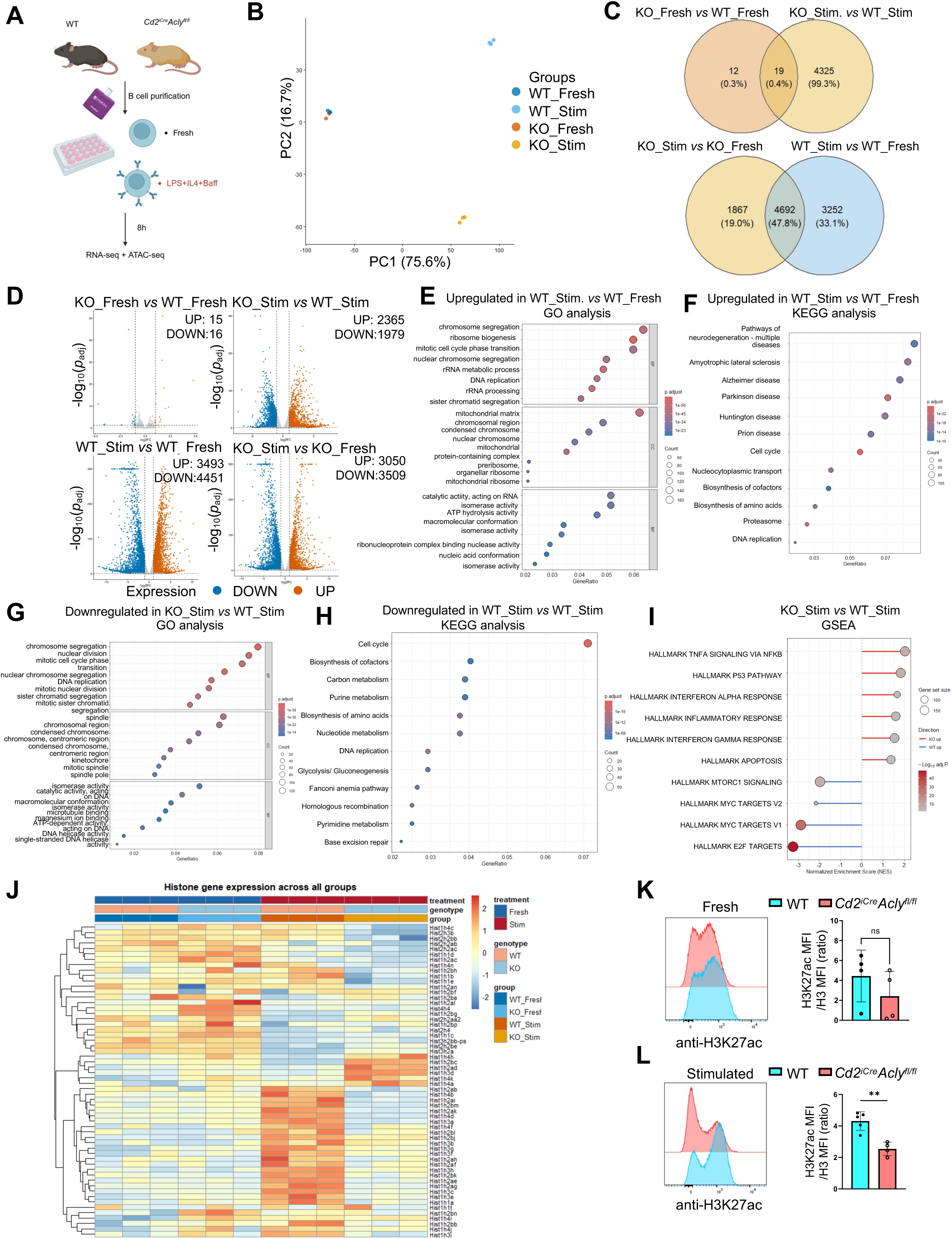
ACLY orchestrates transcriptional reprogramming during B cell activation. (A) Schematic of the experimental design. B cells were isolated from WT and *Cd2^iCre^ Acly*^fl/fl^ mice by magnetic negative sorting and were either maintained in the fresh state or stimulated with LPS, IL-4 and BAFF for 8 h before RNA-seq and ATAC-seq analysis. (B) Principal component analysis (PCA) of transcriptomic profiles across the four experimental groups. WT_Fresh, fresh WT B cells; WT_Stim, stimulated WT B cells; KO_Fresh, fresh Acly-deficient (KO) B cells; KO_Stim, stimulated KO B cells. (C) Venn diagrams showing overlapping differentially expressed genes (DEGs) between KO and WT B cells under fresh and stimulated conditions (top), and between stimulated and fresh states within each genotype (bottom). DEGs were defined using a threshold of adjusted *P* < 0.05 and |log_2_(FC)| > 1. (D) Volcano plots of DEGs in the indicated comparisons: KO_Fresh versus WT_Fresh (top left), KO_Stim versus WT_Stim (top right), WT_Stim versus WT_Fresh (bottom left), and KO_Stim versus KO_Fresh (bottom right). Red and blue dots indicate significantly upregulated and downregulated DEGs, respectively (adjusted *P* < 0.05 and |log_2_FC| > 1). (E, F) Gene Ontology (GO) (E) and KEGG pathway (F) enrichment analyses of genes upregulated in WT_Stim versus WT_Fresh. (G, H) GO (G) and KEGG pathway (H) enrichment analyses of genes downregulated in KO_Stim versus WT_Stim. (I) Gene set enrichment analysis (GSEA) of transcriptomic alterations in KO_Stim versus WT_Stim. (J) Heatmap showing expression patterns of histone-related genes across the indicated groups. (K, L) The expression of H3K27ac (normalized against total H3) on fresh (K) and LPS/IL4/BAFF-activated (L) WT and KO B cells.

Furthermore, we identified a global reduction of histone-associated gene expression in ACLY-deficient cells specifically under stimulated conditions (Fig. 3J). Consistent with this observation, flow cytometry showed no significant difference in the acetylated H3K27 (relative to total H3) between WT and ACLY-deficient cells under fresh conditions, whereas a significant decrease was observed in ACLY-deficient cells upon stimulation (Fig. 3K and L). These results suggest that ACLY is required for activation-induced histone modification.

### ACLY-dependent chromatin accessibility sustains activation-induced regulatory networks

To characterize chromatin accessibility dynamics across conditions, we analyzed differential accessible (DA) peaks among the four experimental groups. KO_Fresh and WT_Fresh exhibited only limited differences in chromatin accessibility (Fig. 4A). In contrast, stimulation triggered extensive chromatin remodeling in WT cells, characterized by widespread gains in accessibility (∼9,000 gained vs <1,000 lost peaks) in WT_Stim relative to WT_Fresh. This activation-associated response was markedly attenuated in ACLY-deficient cells, which instead displayed predominantly losses of accessibility following stimulation (∼11,000 lost vs ∼4,000 gained peaks). Consistently, direct comparison of KO_Stim and WT_Stim revealed a global loss of chromatin accessibility in ACLY-deficient cells (∼8,000 lost vs ∼500 gained peaks) (Fig. 4A). Moreover, transcriptional start site (TSS)-centered accessibility profiling further confirmed a global reduction in chromatin accessibility in KO_Stim compared with WT_Stim (Fig. 4B). These observations contrasted with the largely comparable magnitude of stimulation induced transcriptional alterations in ACLY-deficient cells (Fig. 3D), suggesting that ACLY preferentially modulates chromatin accessibility over transcription.

**Figure 4.**
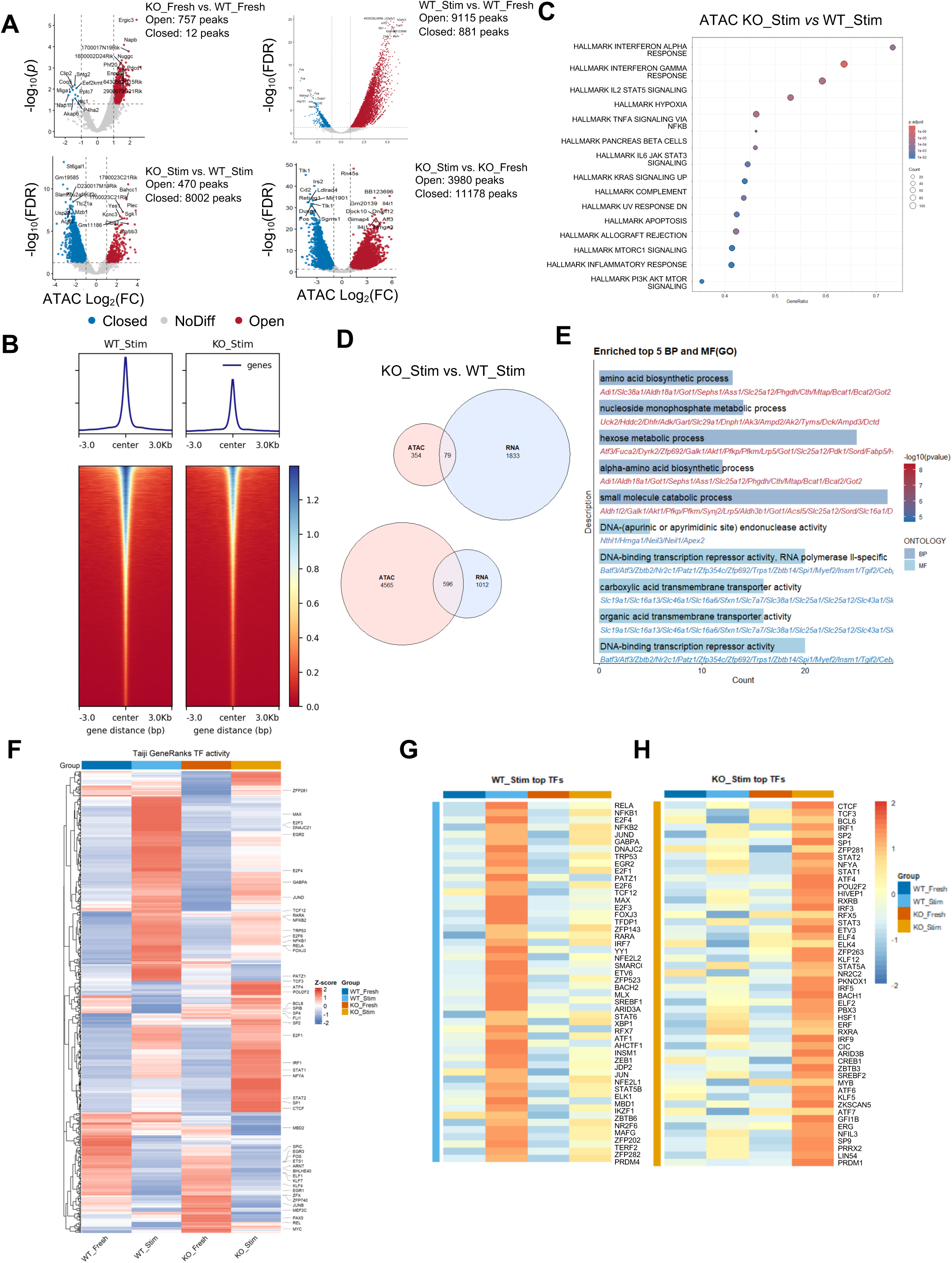
Integrated RNAseq–ATACseq Analyses Reveal ACLY-dependent epigenetic programing following B cell activation. (A) Volcano plots of differentially accessible peaks identified by ATAC-seq in the indicated comparisons: fresh ACLY-deficient (KO) B cells (KO_Fresh) versus fresh WT B cells (WT_Fresh) (top left), stimulated WT B cells (WT_Stim) versus WT_Fresh (top right), stimulated KO B cells (KO_Stim) versus WT_Stim (bottom left), and KO_Stim versus KO_Fresh (bottom right). Red and blue dots denote regions with significantly increased or decreased chromatin accessibility, respectively. Significance thresholds for each comparison are indicated in the corresponding panels. (B) Heatmap of chromatin accessibility surrounding transcription start sites (TSSs) across the indicated groups. (C) GSEA showing the top three significantly enriched pathways in KO_Stim versus WT_Stim. (D) Venn diagrams showing overlap between genes associated with increased chromatin accessibility and upregulated transcription (top), or decreased chromatin accessibility and downregulated transcription (bottom), in KO_Stim versus WT_Stim. DEGs and DAR-associated genes were defined as features with adjusted *P* < 0.05 and |log_2_FC| > 1. (E) GO enrichment analysis of biological process and molecular function categories for the 596 genes exhibiting concordant reductions in chromatin accessibility and transcriptional expression. Top five enriched terms are shown. (F) *Z*-score–normalized PageRank heatmap showing TF activity across WT_Fresh, WT_Stim, KO_Fresh, and KO_Stim. The top 50 TFs ranked by mean rank score across all conditions are annotated. (G, H) Heatmaps showing the top TFs assigned to WT_Stim (G) and KO_Stim (H) condition based on their highest PageRank scores.

To define the functional programs associated with these accessibility changes, we performed pathway enrichment analysis across all differential accessibility comparisons (Fig. 4C and Supplementary Fig. 6A-C). GSEA revealed that WT B cell activation induced broad chromatin opening programs supporting inflammatory and cytokine signaling, including interferon, IL6, IL2, and NFκB (WT_Stim vs WT_Fresh, Supplementary Fig. 6B). In the absence of ACLY, core activation pathways such as NFκB and IL2-Stat5 remained accessible, but key modules, including interferon and IL6-Stat3 signaling, were diminished. In addition, pathways associated with metabolic and biosynthetic functions, including fatty acid metabolism, peroxisome, bile acid metabolism and protein secretion, were suppressed (KO Stim vs KO Fresh, Supplementary Fig. 6C). Importantly, direct comparison of KO_Stim and WT_Stim revealed a uniformly suppressed landscape, with multiple activation-associated pathways–including interferon responses, NFκB, IL2-Stat5, IL6-Stat3, inflammatory response, and PI3K-AKT-mTOR signaling–significantly reduced (Fig. 4C). Collectively, these analyses indicate that ACLY-deficient B cells retain the ability to initiate certain aspects of the activation programs but fail to fully establish or sustain canonical activation-associated chromatin accessibility.

To systematically link transcriptional and chromatin accessibility changes, we integrated RNA-seq and ATAC-seq datasets. Although DEGs were relatively balanced between upregulated and downregulated genes, DA regions were predominantly decreased in KO_Stim, indicating that loss of chromatin accessibility is a dominant feature of ACLY deficiency (Fig. 4D). Focusing on 596 commonly downregulated genes associated with decreased chromatin accessibility, GO analysis revealed significant enrichment in metabolic processes, including amino acid biosynthesis, nucleoside monophosphate metabolism, hexose metabolism, and small molecule catabolic processes (Fig. 4E). Molecular function analysis further highlighted enrichment in DNA-binding transcriptional repressor activity, DNA endonuclease activity, and transmembrane transporter activity. These results indicate that ACLY-dependent chromatin remodeling is tightly coupled to transcriptional regulation of central metabolic and biosynthetic pathways.

To further dissect the regulatory mechanisms underlying these changes, we performed transcription factor (TF) activity analysis using Taiji, which infers TF activity by integrating RNA-seq and ATAC-seq into a PageRank-based gene regulatory network model^16^. Global TF activity profiling revealed distinct clustering across the four experimental conditions (Fig. 4F), indicating extensive remodeling of regulatory networks. Condition-specific TFs, defined by assigning each TF to the condition in which it exhibited maximal PageRank activity, delineated distinct regulatory programs in WT and ACLY-deficient B cells upon stimulation (Fig. 4G,H). KO_Stim displayed a unique TF activity landscape, characterized by altered activation of specific TF subsets compared to WT_Stim, including reduced NFκB, E2F family TFs, and increased IRF1, Stat1/2 activities. Of note, KO_Stim had increased TF activity of CTCF, which regulates cell-cycle arrest and cell death following B cell activation, was consistent with the reduced proliferation and increased apoptosis observed in ACLY-deficient cells^17^. Together, these data support a model in which ACLY couples metabolic state to chromatin accessibility and transcriptional regulation during B cell activation.

### B cell–intrinsic ACLY is required for germinal center formation and antigen-specific humoral immunity *in vivo*

To determine how B cell intrinsic ACLY deficiency could affect humoral immunity *in vivo*, we generated mixed BM chimeras in which BM cells from either WT or *Cd2*^iCre^*Acly*^fl/fl^ mice were mixed with those from μMT mice at 1:4 ratio, and transferred into lethally irradiated congenic CD45.1^+^ mice. In these chimera mice, ACLY deletion was largely restricted to B cells^14^. We immunized the chimera mice with NP-OVA/alum 8 weeks after BM reconstitution. Flow cytometric analysis revealed a marked reduction in Bcl6⁺B220⁺ cells in μMT:*Cd2*^iCre^*Acly*^fl/fl^ chimeras compared with μMT:WT controls (Fig. 5A), suggesting an impaired induction of the GC transcriptional program. Consistently, both the frequency and absolute number of GL7⁺Fas⁺ GC B cells were significantly reduced in the absence of B cell–intrinsic ACLY (Fig. 5B). Analysis of GC compartmentalization further demonstrated a decreased dark zone (DZ) to light zone (LZ) ratio, defined by CXCR4^hi^CD86^lo^ versus CXCR4^lo^CD86^hi^ GC B cells^18^ (Fig. 5C), consistent with disrupted GC organization and dynamics. Moreover, μMT:*Cd2*^iCre^*Acly*^fl/fl^ chimeras exhibited a significant reduction in splenic CD138⁺ plasmablasts (Fig. 5D). Moreover, the frequency of IgG1⁺B220⁺ class-switched B cells was markedly decreased (Fig. 5E). Consistent with these cellular defects, serum ELISA analysis revealed significantly reduced NP23-specific antibody titers, including IgM, IgG1, IgG2b, and IgG3, whereas IgG2c levels were not significantly altered, in μMT:*Cd2*^iCre^*Acly*^fl/fl^ chimeras compared with μMT:WT chimeras (Fig. 5F). Together, these data establish that B cell–intrinsic ACLY is required for efficient GC formation, plasmablast generation, and antigen-specific antibody production following immunization.

**Figure 5.**
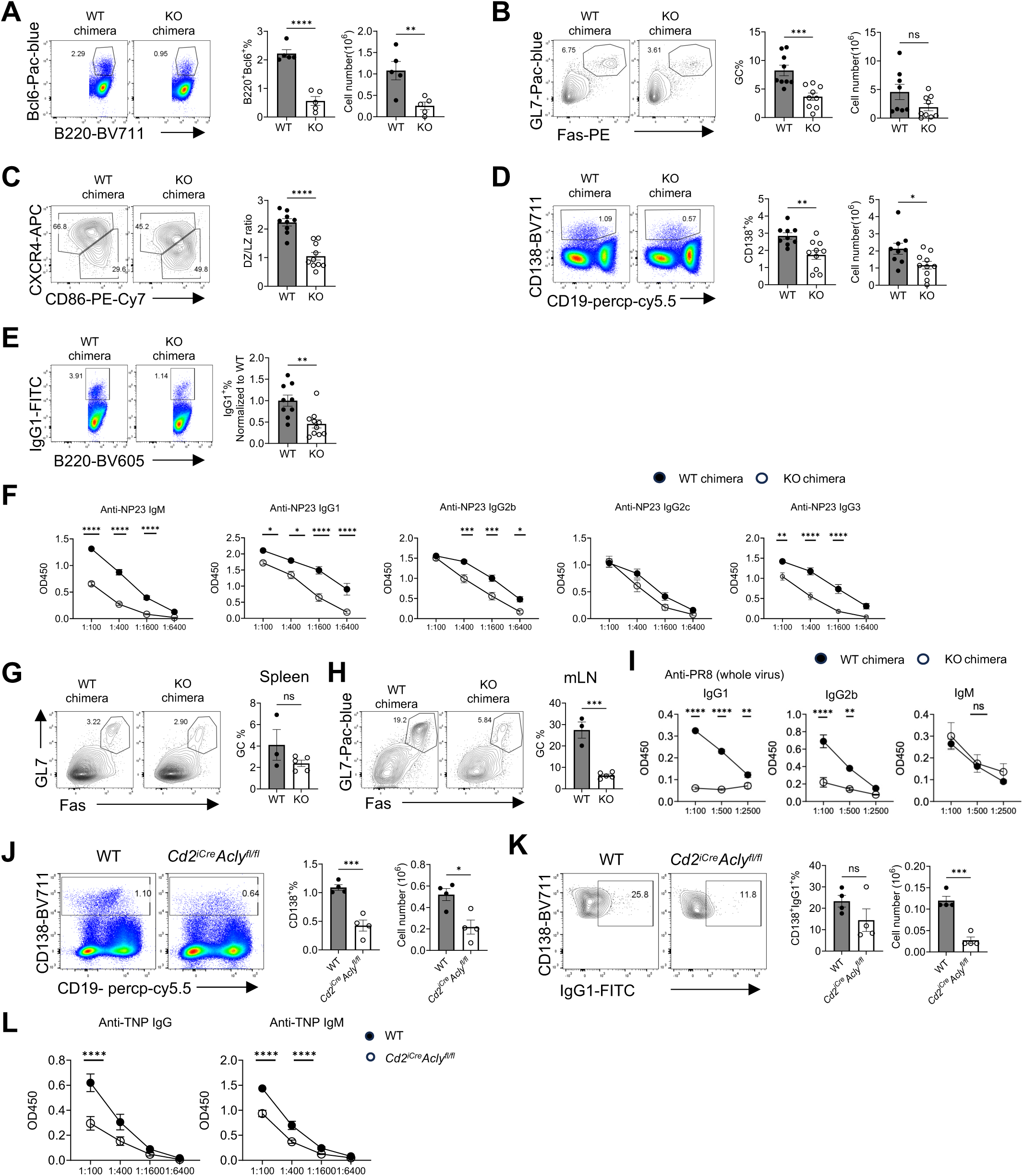
B cell-intrinsic ACLY supports humoral immunity *in vivo*. (A-E) μMT:WT chimera and μMT:*Cd2*^iCre^*Acly*^fl/fl^ chimera mice were immunized with NP-OVA and analyzed after 14 days. Left, flow cytometry of Bcl6 and B220 expression on splenocytes (A), GL7 and Fas expression on B cell (B), CXCR4 and CD86 expression on GC B cell (C), CD19 and CD138 expression on splenocytes (D), and IgG1 and B220 expression on B cell (E). Right, the frequencies and cell number of Bcl6^+^B220^+^ (A), GC B cell (B), CD138^+^ plasmablasts (D), the ratio of DZ (CXCR4^hi^ CD86^low^) and LZ (CXCR4^low^ CD86^hi^) (C), and the frequencies of IgG1^+^B220^+^ (E). (F) Serum anti-NP23 immunoglobulins from NP-OVA immunized mice after 14 days were measured by ELISA. (G-I) μMT:WT chimera and μMT:*Cd2*^iCre^*Acly*^fl/fl^ chimera mice were infected with H1N1 PR8 influenza strain intranasally and were analyzed 11 days after infection. Left, flow cytometry analysis of GL7 and Fas expression on B cell from spleen (G) and mediastinal LN (mLN) (H). Right, the frequency and cell number of GC B cell from spleen (G) and mLN (H). (I) PR8 influenza-specific IgG1, IgG2b and IgM in sera were measured using ELISA (whole virus lysate was used as antigens). (J, K) WT and *Cd2^iCre^ Acly*^fl/fl^ mice were immunized with TNP-LPS and analyzed after 14 days. Left, flow cytometry of CD138^+^ expression (J), and expression of IgG1 on CD138^+^ cells (K) from splenocytes. Right, frequency and cell number of CD138^+^ plasmablasts (J), and IgG1^+^CD138^+^ cell populations (K). (L) Anti-TNP IgG and IgM in sera measured by ELISA. *p* values were calculated using Student t-test or two-way ANOVA. ns, not significant, *p < 0.05, **p < 0.01, ***p < 0.001, ****p < 0.0001. Error bars represent SEM.

To assess whether ACLY is similarly required during antiviral immune responses, μMT:WT and μMT:*Cd2*^iCre^*Acly*^fl/fl^ chimeras were infected with influenza virus (H1N1 PR8 strain) and analyzed 11 days post-infection. μMT:*Cd2*^iCre^*Acly*^fl/fl^ chimeras exhibited a pronounced decrease in the frequency of GC B cells in the mediastinal lymph nodes (mLN), but not in the spleen (Fig. 5G and H). By contrast, plasmablast responses were largely preserved, as no significant differences were detected in the frequency of CD138⁺ cells between μMT:WT and μMT:*Cd2*^iCre^*Acly*^fl/fl^ chimeras in either the spleen or mLN (Supplementary Fig. 4A). Similarly, the frequency of PD-1⁺CXCR5^hi^ T follicular helper (Tfh) cells was not significantly affected by ACLY deletion (Supplementary Fig. 4B). Despite the preservation of plasmablast and Tfh compartments, serum ELISA analysis revealed significantly reduced levels of influenza virus–specific antibodies, including IgG1, and IgG2b, whereas IgM titers were not significantly altered in μMT:*Cd2*^iCre^*Acly*^fl/fl^ chimeras (Fig. 5I). Thus, these findings demonstrate that B cell-intrinsic ACLY is required for effective GC B cell expansion and optimal antiviral humoral immunity.

We next sought to determine if ACLY is required for T cell–independent responses. WT and *Cd2*^iCre^*Acly*^fl/fl^ mice were immunized with T cell–independent antigen TNP-LPS and analyzed after 14 days. *Cd2*^iCre^*Acly*^fl/fl^ mice had significantly reduced frequency and absolute number of CD138⁺ plasmablasts as well as IgG1 expression on the plasmablasts (Fig. 5J and K). Consistent with these cellular changes, serum anti-TNP IgG and IgM titers also reduced significantly in *Cd2*^iCre^*Acly*^fl/fl^ mice (Fig. 5L). Together, these data indicate that ACLY is indispensable for humoral immune responses to both T-dependent and T-independent antigen stimulation.

### ACLY sustains antibody-secreting cell output after B cell activation

So far, our results demonstrate that ACLY supports early B cell activation and survival. To determine whether ACLY is required after B cell activation, we generated *Aicda*^Cre^*Acly*^fl/fl^ mice to ablate ACLY post B cell activation. After immunization with NP-OVA/alum, we did not observe significant differences in terms of frequency and absolute number of GL7⁺Fas⁺ GC B cells between WT and *Aicda*^Cre^*Acly*^fl/fl^ mice (Fig. 6A). Analysis of GC architecture revealed no significant difference in the dark zone (DZ) to light zone (LZ) ratio between genotypes (Fig. 6B). These data suggest that ACLY might be dispensable for GC B cell differentiation post B cell activation. Moreover, *Aicda*^Cre^*Acly*^fl/fl^ mice exhibited a significant reduction in the frequency, but not the absolute number, of splenic CD138⁺ and CD19⁻CD138⁺ plasmablasts (Fig. 6C). By contrast, bone marrow CD138⁺TACI⁺ plasma cells were significantly reduced in *Aicda*^Cre^*Acly*^fl/fl^ mice (Fig. 6D). Consistent with the reduction of antibody producing cells, ELISA analysis revealed significantly reduced serum titers of NP-specific antibodies in *Aicda*^Cre^*Acly*^fl/fl^ mice compared with WT controls (Fig. 6E). Specifically, anti-NP23 IgG1, IgG2b, and IgM levels were modestly decreased, whereas anti-NP2 responses showed more robust and significant reductions in IgG1 and IgG2b but not IgM, suggesting impaired class switching and affinity maturation. Thus, after B cell activation, ACLY is critically required for the generation of antibody producing cells and optimal antibody production, but not for GC formation.

**Figure 6.**
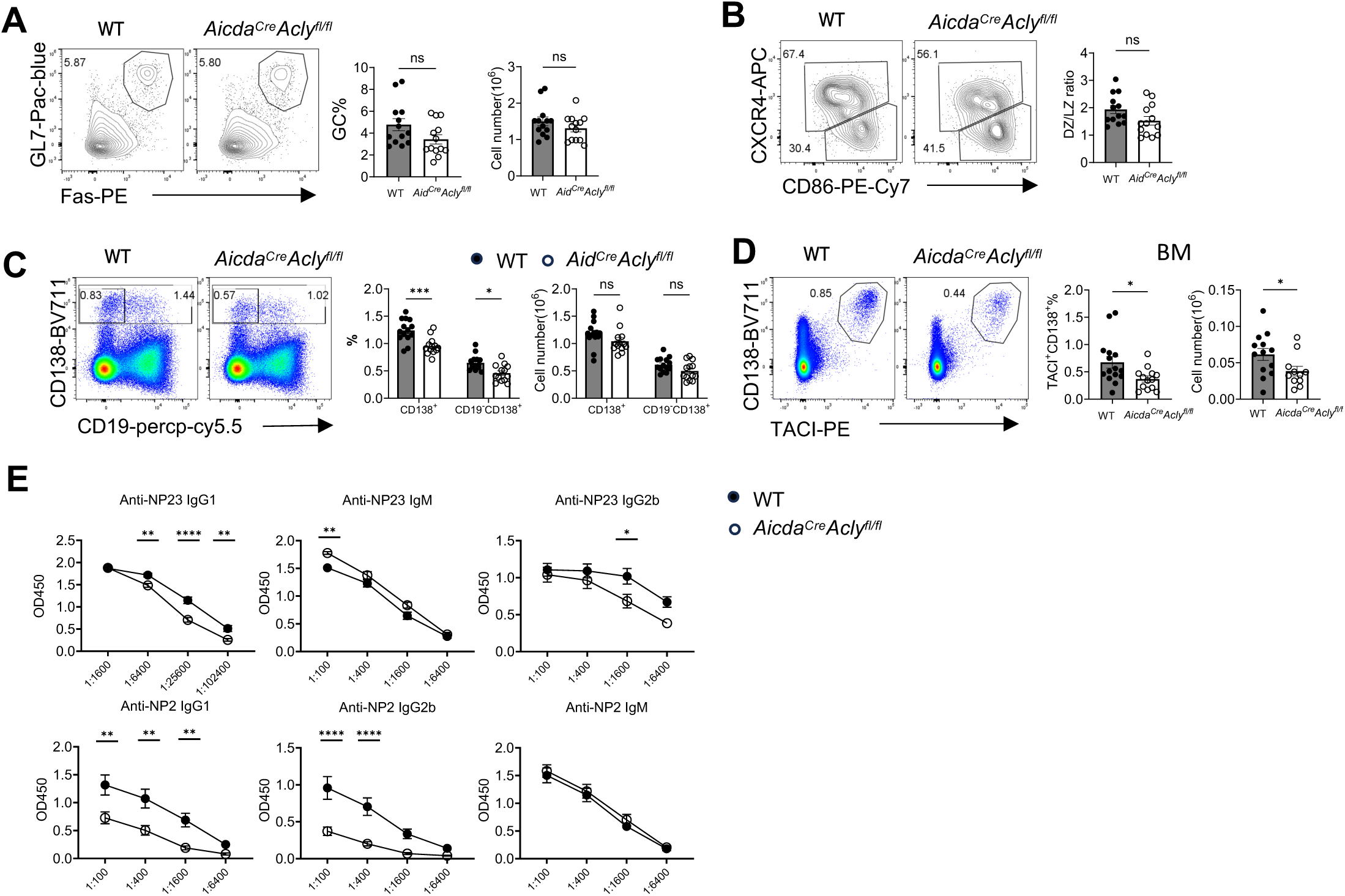
ACLY supports humoral immunity post B cell activation. (A-D) WT and *Aicda^Cre^Acly*^fl/fl^ mice were immunized with NP-OVA/alum and analyzed after 14 days. Left, flow cytometry of GL7 and Fas expression on B cell (A), CXCR4 and CD86 expression on GC B cell (B), CD19 and CD138 expression on splenocytes (C), and CD138 and TACI expression on bone marrow cells (D). Right, the frequencies and cell number of GC B cell (A), CD138^+^ and CD19^-^CD138^+^ cell populations(C), CD138^+^TACI^+^ (D), and the ratio of DZ and LZ (B). (E) ELISA test of serum anti-NP23 and anti-NP2 immunoglobulins from NP-OVA immunized mice presented as absorbance at 450 nM (A_450_). *p* values were calculated using Student t-test or two-way ANOVA. ns, not significant, *p < 0.05, **p < 0.01, ***p < 0.001, ****p < 0.0001. Error bars represent SEM.

To determine whether ACLY contributes to pathological humoral responses, we employed the bm12-induced autoimmune model, in which alloreactive donor CD4^+^ T cells drive sustained germinal center (GC) activation and autoantibody production from host B cells^19^. We transferred splenocytes from bm12 mice to WT and *Aicda*^Cre^*Acly*^fl/fl^ mice, which were analyzed six weeks later (Fig. 7A). Flow cytometric analysis showed that the frequency and absolute number of Bcl6⁺B220⁺ cells and GL7⁺Fas⁺ GC B cells were comparable between WT and *Aicda*^Cre^*Acly*^fl/fl^ mice (Fig. 7B,C), indicating that ACLY deletion post B cell activation does not overtly affect GC establishment under chronic autoimmune stimulation. In contrast, GC organization was altered, as evidenced by a significantly reduced dark zone (DZ) to light zone (LZ) ratio in *Aicda*^Cre^*Acly*^fl/fl^ recipients (Fig. 7D). *Aicda*^Cre^*Acly*^fl/fl^ recipients exhibited a significant reduction in both the frequency and absolute number of CD19^int^CD138⁺ plasmablasts (Fig. 7E). Functional assessment of autoantibody production revealed that overall serum levels of anti–double-stranded DNA antibodies were comparable between WT and *Aicda*^Cre^*Acly*^fl/fl^ recipients (Fig. 7F). While anti-histone IgG1 levels were unchanged, anti-histone IgG2b and IgG2c titers were significantly reduced in *Aicda*^Cre^*Acly*^fl/fl^ mice (Fig. 7G). These findings indicate a selective defect in pathogenic IgG subclass production rather than a global reduction of autoreactive antibody responses. Together, these data demonstrate that ACLY is required to sustain GC organization, class-switch diversification, and pathogenic IgG subclass output during chronic immune activation, extending its role beyond protective humoral immunity to the regulation of autoimmune antibody responses.

**Figure 7.**
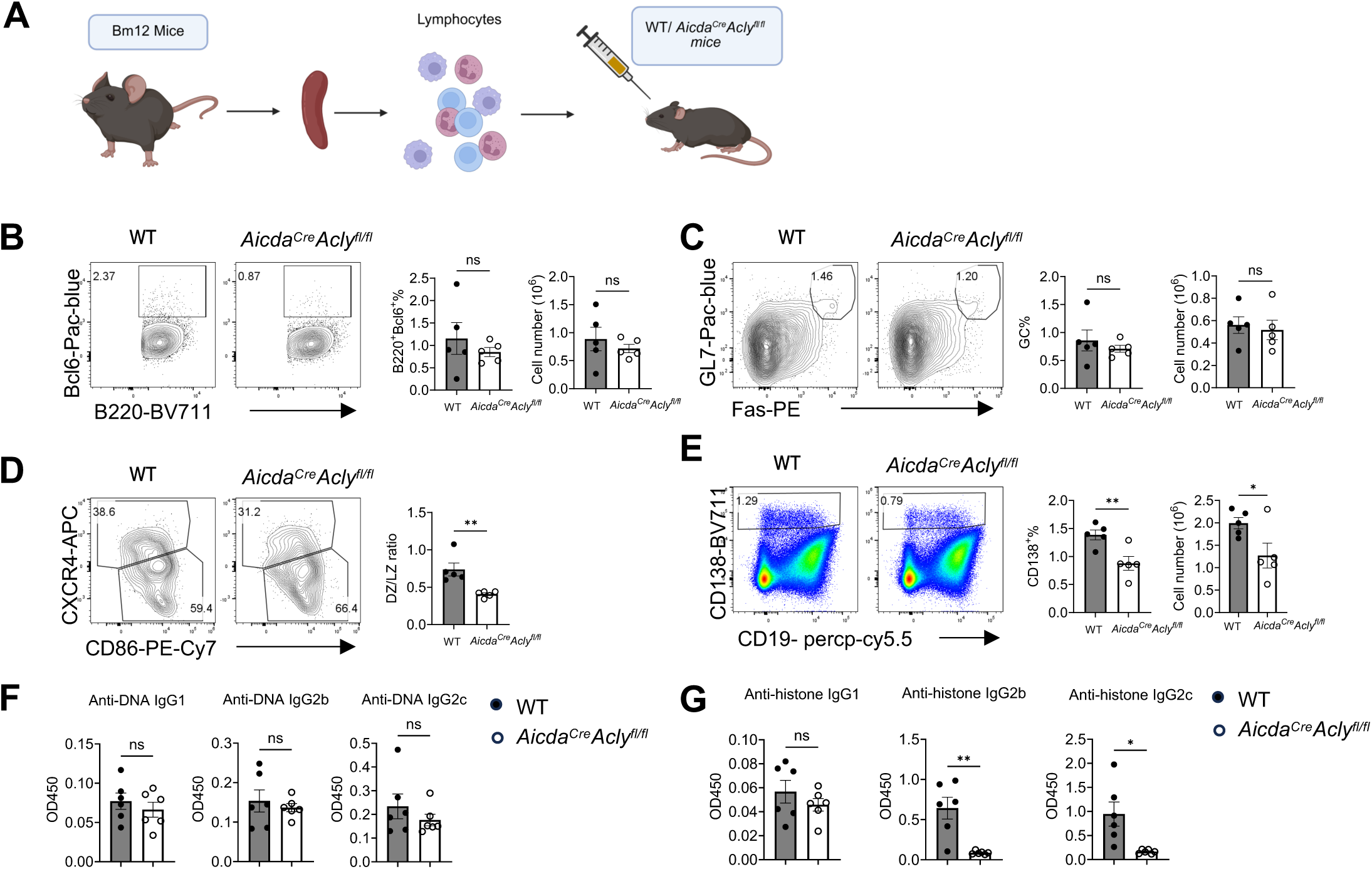
ACLY supports autoreactive B cell response. (A) The schematic diagram of Bm12 induced autoimmune mouse model on *Aicda^Cre^Acly^fl/fl^*mice. About 30 million lymphocytes from Bm12 mice were transferred into each WT and *Aicda^Cre^Acly^fl/fl^* mouse. And the mice were analyzed after 6 weeks. (B-E) Left, flow cytometry of Bcl6 and B220 expression on splenocytes (B), GL7 and Fas expression on B cell (C), CXCR4 and CD86 expression on GC B cell (D), CD19 and CD138 expression on splenocytes (E). Right, the frequencies and cell number of Bcl6^+^B220^+^ (B), GC B cell (C), the ratio of DZ and LZ (D), CD19^int^CD138^+^ plasmablasts (E). (F, G) ELISA test of serum anti-DNA (F) and anti-histone immunoglobulins (G). *p* values were calculated using Student t-test (B-G). ns, not significant, *p < 0.05, **p < 0.01. Error bars represent SEM.

## Discussion

Although there have been tremendous advances in our understanding of signaling events and metabolic programs downstream of BCR and TLR, the epigenetic program during early B cell activation remains largely unknown. Recent studies have established ACLY as a key regulator of epigenetic programs in effector T cell differentiation and T cell exhaustion. However, ACLY deficiency does not affect T cell activation or survival, suggesting that ACLY dependent metabolic-epigenetic program is largely dispensable for T cell activation. Our study contrasts sharply with the published studies and demonstrates that TLR/BCR elicited B cell activation and survival critically depend on ACLY mediated metabolic and epigenetic programing.

Normal B cell activation is accompanied by increased uptake of glucose^20–22^, which is either metabolized through glycolysis^23^, or fed into biosynthetic pathways^24,25^. ACLY is known to convert glucose derived citrate into acetyl-CoA in the cytosol or nucleus, thereby supplying the essential precursor for all lipid biosynthesis and protein acetylation. Consistent with our previous findings that fatty acid availability is critically required for B cell activation and survival, we find here that ACLY is dispensable for B cell development and homeostasis but becomes functionally important upon activation, when its expression is greatly enhanced. Importantly, the dependence of ACLY for B cell proliferation is selective. Whereas TLR/BCR induced proliferation is dependent on ACLY, lymphopenia induced homeostatic proliferation is not, consistent with normal B cell homeostasis in *Cd2*^iCre^*Acly*^fl/fl^ mice. Current model posits that BAFF and tonic BCR signaling support B cell homeostatic proliferation^26,27^. Thus, our data suggests that BAFF and tonic BCR signaling may induce a unique metabolic, transcriptional and epigenetic program that is completely ACLY-independent. This question awaits future investigation.

To define ACLY’s role, we integrated RNA-seq and ATAC-seq across four comparisons. At baseline, ACLY deficiency had minimal impact on both gene expression and chromatin accessibility, although enrichment of inflammatory and stress-associated pathways suggests subtle epigenetic priming in resting cells. Upon stimulation, WT B cells underwent extensive chromatin remodeling accompanied by robust transcriptional changes, with accessibility enriched at canonical activation pathways, including NFκB, interferon, and STAT signaling, as well as cell cycle–associated programs.

In contrast, ACLY-deficient B cells exhibited a markedly impaired epigenetic response to stimulation, characterized by a global reduction in chromatin accessibility. Despite this, the overall magnitude of transcriptional responses remained broadly comparable to WT cells. Pathway analysis revealed that ACLY deficiency broadly attenuated accessibility across canonical activation-associated programs, including interferon, NFkB, IL2-Stat5, IL6-Stat3, inflammatory response, and PI3K-AKT-mTOR signaling, with no evidence of compensatory pathway enrichment. These findings indicate that ACLY is required to sustain, rather than merely initiate, activation-associated chromatin remodeling.

The divergence between chromatin accessibility and transcription output suggests that ACLY primarily regulates the epigenetic potential of activated B cells. Whereas ATAC-seq captures accessibility at regulatory elements, RNA-seq reflects the integrated outcome of downstream processes, including transcription factor activity and signaling feedback. As such, widespread attenuation of chromatin accessibility can coexist with partially preserved transcriptional responses. Therefore, these findings identify ACLY as a key metabolic regulator of activation-induced chromatin remodeling. By supporting acetyl-CoA production, ACLY enables the establishment of a chromatin landscape that sustains canonical immune signaling during B cell activation.

The physiological importance of B cell–intrinsic ACLY is underscored by the marked reduction in antigen-specific antibody titers observed in both peptide immunization and influenza infection models. ACLY deficiency reduces both germinal center (GC) B cells and plasmablasts following immunization, but selectively impairs GC formation without affecting plasmablast generation during influenza infection, revealing a context-dependent requirement for ACLY in plasmablast differentiation.

To delineate stage-specific functions, we employed an Aicda-Cre–mediated *Acly* floxed model to delete ACLY after B cell activation, thereby bypassing its role in early activation. Under these conditions, ACLY is required for plasmablast generation but is dispensable for GC formation. Consistent with this, post-activation deletion of ACLY significantly reduces plasmablast responses and autoantibody production in a chronic graft-versus-host disease (cGvHD) model, supporting an important and distinct function of ACLY post B cell activation. These results reminiscent of a previous study on the stage specific functions of LDHA in humoral immunity^23^.

Together, these findings establish that ACLY plays distinct and stage-specific roles in B cell responses, being required both during early activation and for subsequent plasmablast differentiation. These results position ACLY as a likely key metabolic regulator of humoral immunity and autoimmunity. The molecular mechanisms linking ACLY activity to B cell fate decisions following activation remain to be defined.

## Materials and Methods

### Mice

*Cd2^iCre^Acly*^fl/fl^ mice and *Aicda^Cre^Acly*^fl/fl^ mice were generated by crossing *Acly2*^fl/fl^ mice (*Acly^tm1.1Welk^*/Mmjax, from MMRRC) with *Cd2-iCre* transgenic mice and *Aicda-Cre* mice, respectively, purchased from Jackson Laboratory. C57BL/6, B6.CD45.1 (B6.SJL-*Ptprc^a^ Pepc^b^*/BoyJ), and *Rag1*^−/–^ mice, B6.129S2-*Ighm^tm1Cgn^*/J (μMT) mice, B6(C)-*H2-Ab1^bm12^*/KhEgJ (Bm12) mice were purchased from the Jackson Laboratory.

In the *Cd2^iCre^Acly*^fl/fl^-μMT bone marrow (BM) chimera mouse model, BM cells from *Cd2^iCre^Acly*^fl/fl^ (knockout, KO) and *Acly*^fl/fl^ (wild type, WT) mice were harvested. These were then mixed separately with BM cells from μMT mice at a ratio of 1:4. T cells were depleted from all BM cells using Miltenyi anti-CD4 and anti-CD8 microbeads. The chimeric mice were created by transferring BM cells into lethally irradiated B6.CD45.1 mice. The immune system was subsequently reconstituted after 8 weeks.

*Cd2^iCre^Acly^fl/fl^-Rag1* chimera mouse model were generated by transferring B cells isolated from *Cd2^iCre^Acly*^fl/fl^ or *Acly*^fl/fl^ mice, mixed with B cells from B6.CD45.1 mice at a ratio of 1:1 into *Rag1^−/–^* mice. All B cells were labeled with Celltrace violet (CTV).

Mice were maintained under specific pathogen-free conditions within the Department of Comparative Medicine of Mayo Clinic. All the animal experiments were approved by the Institutional Animal Care and Use Committee (IACUC).

### In vivo immunization model

For NP-OVA/Alum immunization, 100 µg NP-OVA (BioSearch Technologies) and 10% KAL(SO_4_)_2_ (Sigma) dissolved in PBS were mixed at a ratio of 1:1, along with 10 µg LPS (Sigma-Aldrich) at pH 7 for each mouse. For TNP-LPS immunization, 50 µg of TNP-LPS (Biosearch Technologies) was dissolved in 200 µl PBS for each mouse. The mice were immunized via intraperitoneal injection.

### Influenza Virus Infection

For influenza virus infection, influenza A/PR8 strain (60 pfu/mouse, gift from Dr. Jie Sun) were diluted in FBS-free DMEM media on ice and inoculated in anesthetized mice through intranasal route. Sera were collected two weeks after infection for immunoglobulins measurement. The mediastinal lymph nodes and spleens were harvested and analyzed for germinal center B cell and plasmablast formation and follicular helper T differentiation.

### Immune cell purification and culture

Mouse B cells or CD4^+^ T cells were isolated from spleen single cell suspension using Mouse B Cell Isolation Kit (StemCell Technologies). Cells were cultured in RPMI 1640 (Corning) containing 10% FBS (Gibco), 10 mM HEPES (Gibco), Penicillin Streptomycin Glutamine mixed solution (Gibco) and 50 µM 2-Mercaptoethanol (Sigma-Aldrich). B cells were stimulated with 10 µg/mL LPS (Sigma-Aldrich), 10 ng/mL recombinant IL-4 (Tonbo Bioscience) and 20 ng/mL recombinant BAFF (BioLegend), or 10 µg/mL anti-mouse IgM (Jackson Immunoresearch), 10 ng/mL recombinant IL-4 and 20 ng/mL recombinant BAFF. CD4^+^ T cells were activated with plate coated anti-CD3 and anti-CD28 (Bio X Cell). Cells were treated with Oleic Acid-Albumin from bovine serum (Sigma-Aldrich), sodium acetate (fisher chemicals), cholesterol (Sigma Aldrich), or Acetyl coenzyme A (Sigma-Aldrich). Proliferation was measured by dilution of CellTrace Violet (CTV) proliferation dye (Life Technologies).

### Real-time PCR

For mRNA analysis, total mRNA was extracted from mouse B cells by RNeasy Micro kit (QIAGEN) and converted to cDNA by using PrimeScript RT Master Mix (Takara), following the manufacturer’s protocols. This was done in preparation for subsequent real-time PCR analysis by using a Takara Realtime PCR system. The following primer was used, *Acly* primers, forward, 5’-AAGCCTTTGACAGCGGCATCATTC-3’, reverse, 5’-TTGAGGATCTGCACTCGCATGTCT-3’. β-actin expression was used as control.

### Metabolic Assays

The bioenergetic activities were measured using a Seahorse XFe96 Extracellular Flux Analyser following established protocols (Agilent). Briefly, about 300,000 B cells were seeded per well on Cell-Tak (Corning) coated XFe96 plate with fresh XF media (Seahorse XF RPMI medium containing 10 mM glucose, 2 mM L-glutamine, and 1 mM sodium pyruvate, PH 7.4; all reagents from Agilent). Oxygen consumption rates (OCR) were measured in the presence of Oligomycin (1.5 µM; Sigma-Aldrich), FCCP (1.5 µM; Sigma-Aldrich), and Rotenone (1 µM; Sigma-Aldrich)/ Antimycin A (1 µM; Sigma-Aldrich) in Mito Stress assay.

### Flow Cytometry

Single-cell suspension from spleens, peripheral lymph nodes, thymus, bone marrow or Peyer’s patches or mediastinal lymph nodes were prepared in PBS containing 2% (w/v) FBS on ice. For analysis of surface markers, cells were stained in PBS containing 1% (w/v) BSA on ice for 30 min, with BV605-labelled B220 (BioLegend, Cat# 103244), PE-labelled anti-CD43 (BD Biosciences, Cat# 553271), Percp-cy5.5 –labelled anti-CD19 (BioLegend, Cat#), PE/Cy7-labelled anti-CD21 (BioLegend, Cat# 123420), FITC-labelled anti-CD23 (BioLegend, Cat# 101606), Alexa Fluor647-labelled anti-CD95/Fas (BD Biosciences, Cat# 563647), Alexa Fluor 488-labelled anti-GL7 (BioLegend, Cat# 144611), APC-eFluor 780-labelled TCRβ (Life Technologies, Cat# 47-5961-82), BV605-labelled anti-CD4 (BioLegend, Cat#), PE-labelled anti-CD8 (BioLegend, Cat# 100707), Alexa Fluor 488-labelled anti-IgG1 (BioLegend, Cat# 406626), BV711-labelled anti-CD138 (BioLegend, Cat# 142519), APC/Cy7-labelled anti-IgD (BioLegend, Cat# 405715), APC-labelled anti-ICOS (BD Biosciences, Cat# 313510), PE-labelled anti PD-1 (BioLegend, Cat# 12-9969-41), Super Bright 600-labelled anti-CD25 (BioLegend, Cat# 63-0251-82), PE/Cy7-labelled anti-CD24 (BioLegend, Cat# 101822), BV711-labelled anti-CD44 (clone: IM7; BioLegend, Cat# 103057). PE-labelled anti-CD267 (TACI) (BioLegend, Cat# 133403), PE/Cy7-labelled anti-CD69 (Tonbo, Cat# 60-0691-U100), APC-labelled anti-CD71 (Life Technologies, Cat# 17-0711-82), Alexa Fluor 647-labelled anti-CD98 (BioLegend, Cat# 128210), APC-labelled anti-CXCR4 (Life Technologies, Cat# 17-9991-82), PE/Cy7-labelled anti-CD86 (BioLegend, Cat# 105014). Cell viability was stained using the Ghost Dye Violet 510 (Cytek) or 7AAD. CXCR5 was stained with biotinylated anti-CXCR5 (BD Biosciences, Cat# 551960) followed by staining with streptavidin-conjugated PE (Cytek Biosciences, Cat# 50-4317). Transcriptional factors Foxp3 (Cytek Biosciences, Cat# 20-5773-U100), Pacific Blue-labelled anti-Bcl6 (BD Biosciences, Cat# 563363), ATP-Citrate Lyase antibody (ACLY) (Cell signaling, Cat# 4332S), and Acetyl-Histone H3(K27) (Cell signaling, Cat# 8173T) were stained using the Transcription Factor Buffer Set (True-Nuclear, BioLegend). Phosflow staining for AF647 conjugated phosphor-S6 (Ser240/244) (Cell signaling, Cat# 4851S) was performed using Phosflow Fix/Perm kit (BD Biosciences). FITC-Annexin V (BioLegend) and Annexin V Binding Buffer (SouthernBiotech) were utilized for apoptosis staining. MitoTracker Deep Red (Invitrogen) and MitoTracker Green (Invitrogen) were utilized for mitochondria staining. And BODIPY™ 493/503 (Invitrogen) were utilized for neutral lipid staining. Flow cytometry was performed on the Attune NxT (ThermoFisher) cytometer, and analysis was performed using FlowJo software (BD Biosciences).

### Mouse Immunoglobulin Isotyping Panel Detection

To detect the concentrations of mouse immunoglobulin isotypes IgG1, IgG2a/IgG2c, IgG2b, IgG3 and total IgM and IgA in sera, we used LEGENDplex mouse immunoglobulin isotyping panel, according to manufacturer’s instructions (BioLegend).

### ELISA

To measure the levels of NP-specific antibodies in sera, 96-well plates (2596; Costar) were coated with 1 µg/mL NP23-BSA or NP2-BSA (Biosearch Technologies) in PBS and incubated at 4°C overnight. The plates were washed four times using 0.05% Tween 20 in PBS, then blocked with 5% blocking protein (Bio-Rad) for 1.5 h at 37°C and washed four times again. Serum samples, serially diluted with 1% BSA, were incubated in the plates for 1.5 h at 37°C and then washed eight times. Horseradish peroxidase (HRP)-conjugated detection antibodies for total IgG, IgG1, IgG2b, IgG2c, IgG3, and IgM (Bethyl Laboratories) were added and incubated at room temperature for 1h, followed by eight washes. The reaction was developed with tetramethylbenzidine (TMB) substrate and stopped with 2N H_2_SO_4_. The absorbance was read at a wavelength of 450 nm. Similar methods were used to detect antibodies specific to the influenza A/PR8 strain (a gift from Dr. Jie Sun, University of Virginia, USA), TNP-LPS (Biosearch Technologies), anti-DNA (homemade), and anti-histone (Sigma Aldrich) in sera.

### Immunoblotting

For immunoblotting, B cells were isolated from mouse spleen with EasySep Mouse B cell isolation kit (STEMCELL Technologies). Retroviral transduced B cells were sorted on a BD FACS Melody sorter. Cells were lysed in lysis buffer with protease and phosphatase inhibitors (Sigma-Aldrich). The transferred membrane was blocked with TBST (0.1% Tween 20) containing 5% BSA for 1 h at room temperature. The membrane was incubated with primary antibodies overnight including ATP-Citrate Lyase antibody (Cell signaling), AceCS1 (clone: D19C6; Cell signaling), p-Akt (Ser473) (clone: D9E; Cell signaling), p-BLNK (Thr152) (clone: E4P2P; Cell signaling), p-SYK (clone: C87C1; Cell signaling), 4E-BP1 (clone:53H11; Cell signaling), c-Myc (clone: E5Q6W; Cell signaling), p-S6 Ribosomal Protein (Ser235/236) (clone: D57.2.2E; Cell signaling), c-Myc (clone: D84C12; Cell signaling) and anti-β-actin (clone: 13E5; Sigma-Aldrich). Then, the membrane was washed and incubated with the corresponding secondary antibody for subsequent enhanced chemiluminescence exposure.

### RNA isolation, library construction, and sequencing

Total RNA was extracted from freshly isolated B cells and B cells stimulated with LPS, IL-4, and BAFF for 7 h using the RNeasy Micro Kit, according to the manufacturer’s instructions. Following quality assessment, high-quality RNA samples were used for library preparation. RNA sequencing libraries were constructed and sequenced on the MGISEQ-2000 platform (BGI Genomics, Shenzhen, China).

### RNA-seq data processing and analysis

RNA-seq reads were aligned to the mouse reference transcriptome (GRCm38/mm10) using STAR. Gene-level expression was quantified with RSEM (v1.3.1). Differential expression analysis was conducted in R (v4.5.1) using DESeq2 (v1.50.2). Genes with an adjusted *P* value < 0.05 (Benjamini–Hochberg correction) and |log2FC| > 1 were considered significantly differentially expressed. Gene Ontology (GO) biological process enrichment analysis was performed using clusterProfiler (v4.18.4). Principal component analysis (PCA) was conducted based on normalized gene expression values obtained from DESeq2.

### ATAC-seq

ATAC-seq data were processed using the HiChIP pipeline^28^. Paired-end sequencing reads were aligned to the mouse reference genome (mm10) using Burrows-Wheeler Aligner (BWA, v0.5.9). To eliminate ambiguously mapped reads, additional in-house filtering was applied to retain only read pairs in which both ends mapped uniquely to a single genomic location. Reads generated from PCR amplification artifacts were further removed as duplicates using Picard MarkDuplicates (v1.67). Open chromatin regions were identified using the “Model-based Analysis of ChIP-Seq” (MACS2, v2.0.10) peak-calling algorithm. Peaks with a false discovery rate (FDR) cutoff of ≤1% were considered statistically significant and retained for downstream analyses. Genes nearest to identified peaks, along with distances from transcription start sites (TSS), were annotated using HOMER annotatePeaks. Peak enrichment across samples was visualized using normalized tag density profiles generated in 200 bp windows with a 20 bp step size using Bedtools (v2.16.2)^29^ together with in-house Perl scripts. Differential accessibility analysis between conditions was performed using DiffBind (v2.14.0), applying significance thresholds of FDR < 0.05 and absolute fold change >2. Additional visualizations of differential peaks and chromosomal distributions were generated using custom R scripts.

### Transcription factor activity analysis

Transcription factor (TF) activity was inferred using Taiji (v2.0)^16^ by integrating RNA-seq and ATAC-seq data. Briefly, Taiji constructs a regulatory network by linking chromatin accessibility with gene expression and estimates TF activity using a PageRank-based algorithm. The resulting TF activity scores were used for downstream visualization, and selected TFs were displayed as heatmaps to illustrate activity patterns across conditions.

### Untargeted metabolomics

B cells were stimulated with LPS, IL-4, and BAFF for 16 hours. Following activation, cells were washed three times with ice-cold PBS, and cell pellets were collected for metabolomic analysis. Briefly, metabolites were extracted from snap-frozen cell pellets using ice-cold acetonitrile:methanol (1:1) containing ^13^C_6_-phenylalanine as an internal standard. Metabolomic profiling was conducted on an Agilent 6550 quadrupole time-of-flight mass spectrometer coupled to an Agilent 1290 Infinity UHPLC system. Metabolites were profiled in both positive and negative electrospray ionization modes and separated using hydrophilic interaction liquid chromatography and reverse-phase C18 columns, generating four acquisition modes per sample. Raw data processing, feature alignment and peak extraction were performed using Agilent Mass Profiler Professional.

For downstream analysis, features with peak intensity <5,000 in more than 25% of samples were excluded. Data were normalized by constant-sum normalization, log transformed and mean centered using MetaboAnalystR. Principal component analysis (PCA) and hierarchical clustering were performed to evaluate sample distribution and group separation. Differential metabolite abundance between groups was assessed using unpaired two-sided Student’s t-tests with multiple-testing correction. Metabolites with an FDR-adjusted P value ≤0.05 and |fold change| ≥1.5 were considered significantly altered. Pathway enrichment analysis was performed using MetaboAnalyst 6.0.

### Statistical Analysis

Statistical analysis was performed using GraphPad Prism (version 9). Comparisons between two groups were examined using a two-tailed Student’s t-test or two-way ANOVA. Schematic were created by BioRender. A p-value < 0.05 was considered statistically significant.

## Acknowledgement

The authors would like to thank Dr. Jie Sun (University of Virginia) for sharing research materials with us. We thank Suwanlikit Yossawat, Fengxin Gao and Shyang Hong Tan for the assistance of bioinformatics. We thank the manager of Mayo Clinic Metabolomics Core facility, Mai Petterson, for her assistance with the metabolomics data analysis and interpretation.

## Author Contributions

Conceptualization: M.L., Z.Z., and H.Z.; Methodology: M.L., Z.Z., X.Z.,Y.L., Xingxing.Z. A.B., N.K.N., and H.Z.; Validation: M.L., Z.Z., and H.Z.; Data curation: M.L., Z.Z., A.B., N.K.N., and H.Z.; Software: Z.Z., A.B., N.K.N.; Formal analysis: M.L., Z.Z., and H.Z.; Investigation: M.L., Z.Z. and H.Z.; Resources: J.S., H.Z.; Writing–original draft: M.L., Z.Z., and H.Z.; Writing–review and editing: M.L., Z.Z., X.Z., and H.Z.; Visualization: M.L., Z.Z., A.B., N.K.N., and H.Z.; Funding acquisition: H.Z. This work was primarily conducted at Mayo Clinic Rochester.

## Data, code, and material availability

All data and code needed to evaluate and reproduce the results in the paper are present in the paper and/or the supplementary Materials. Sequencing data are available at the NCBI BioProject Sequence Read Archive (SRA) (PRJNA1370617). This study does not generate new reagents.

## Figure Legends

**Supplementary Figure 1.**
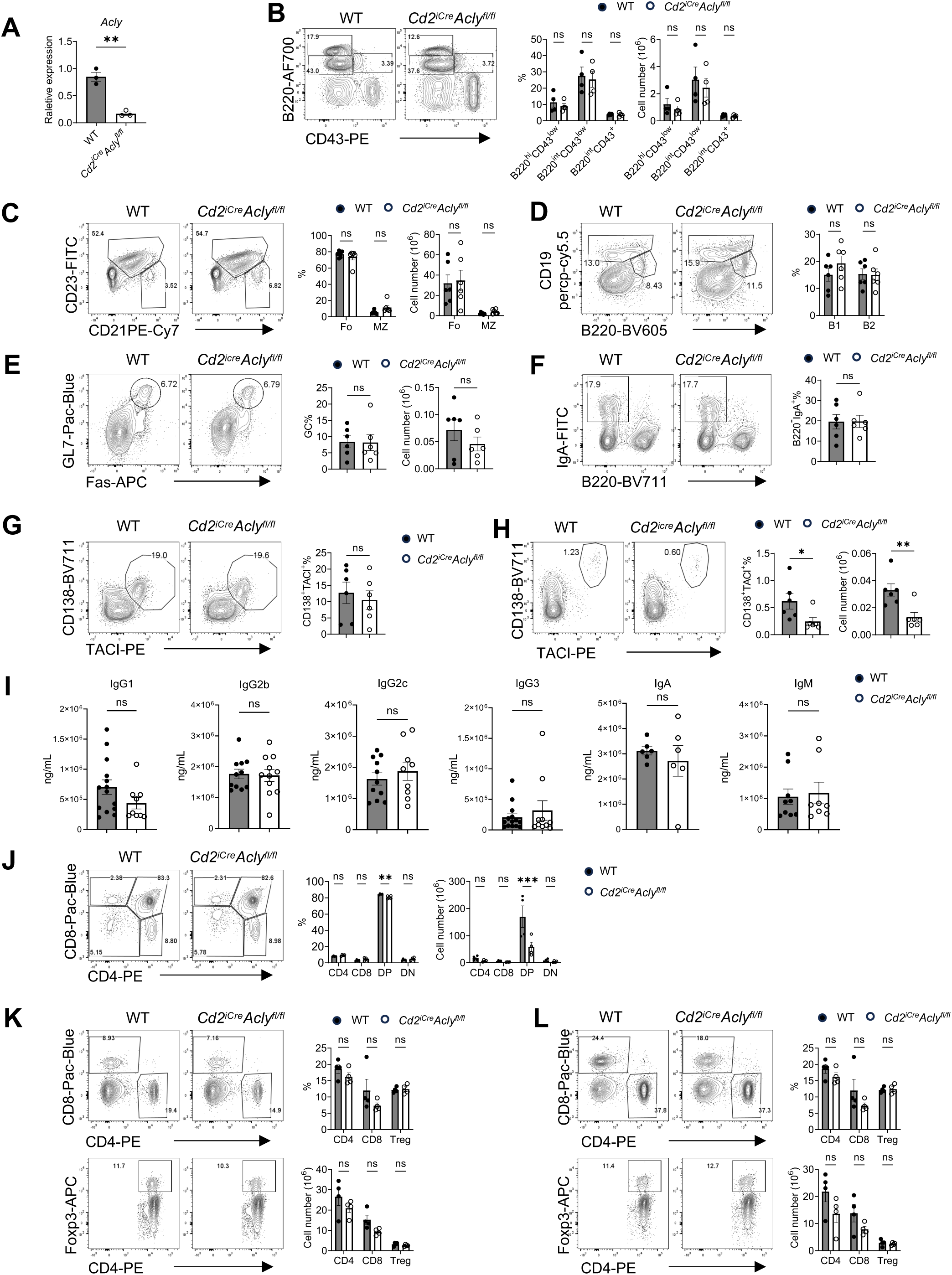
B and T cell development and homeostasis in *Cd2^iCre^Acly*^fl/fl^ mice. (A) Real-time PCR analysis of *Acly* from B cells isolated from WT and *Cd2^iCre^Acly*^fl/fl^ mice’ spleens. (B) Left, the expression of B220 and CD43 of BM cells. Right, the frequency (left) and absolute number (right) of B220^hi^CD43^low^, B220^int^CD43^low^ and B220^int^CD43^+^ cell populations. (C) Left, flow cytometry analysis of Fo (CD21^low^CD23^+^), MZ (CD21^hi^CD23^−^) cells from spleens. Right, the frequency (left) and absolute number (right) of Fo and MZ cell populations. (D) B1 (CD19^hi^B220^low^) and B2 (CD19^int^B220^hi^) cell populations in peritoneal fluid from WT and *Cd2^iCre^Acly*^fl/fl^ mice. (E) Left, flow cytometry analysis of GC (GL7^+^Fas^+^) among TCRβ^-^CD19^+^ population from Peyer’s patches. Right, the frequency (left) and absolute number (right) of GC populations. (F) Left, the expression of B220 and IgA on colon lamina propria (LPL) cells. Right, the frequency of B220^-^IgA^+^. (G) Left, the expression of CD138 and TACI on colon LPL cells. Right, the frequency (left) and absolute number (right) of CD138^+^TACI ^+^. (H) Left, the expression of CD138 and TACI on BM cells. Right, the frequency (left) and absolute number (right) of CD138^+^TACI ^+^. (I) The baseline levels of IgG1, IgG2a, IgG2b, IgG3, IgA and IgM in mouse sera collected from WT and *Cd2^iCre^Acly*^fl/fl^ mice. (J) Left, the expression of CD4 and CD8 on thymus lymphocytes analyzed by flow cytometry. Right, the frequency (left) and absolute number (right) of DN (CD4^-^CD8^-^), DP (CD4^+^CD8^+^), CD4 SP (CD4^+^CD8^-^) and CD8 SP (CD4^-^CD8^+^) cell populations. (K-L) Left, flow cytometry analysis of CD4, CD8 and Treg cells (B220^-^CD4^+^Foxp3^+^) from spleens (L) or pLN (M). Right, the frequency (left) and absolute number (right) of CD4, CD8 and Treg cells. *p* values were calculated using Student’s t test. ns, not significant, *p < 0.05, **p < 0.01, ***p < 0.001. Data are representative of at least three independent experiments. Error bars represent SEM.

**Supplementary Figure 2.**
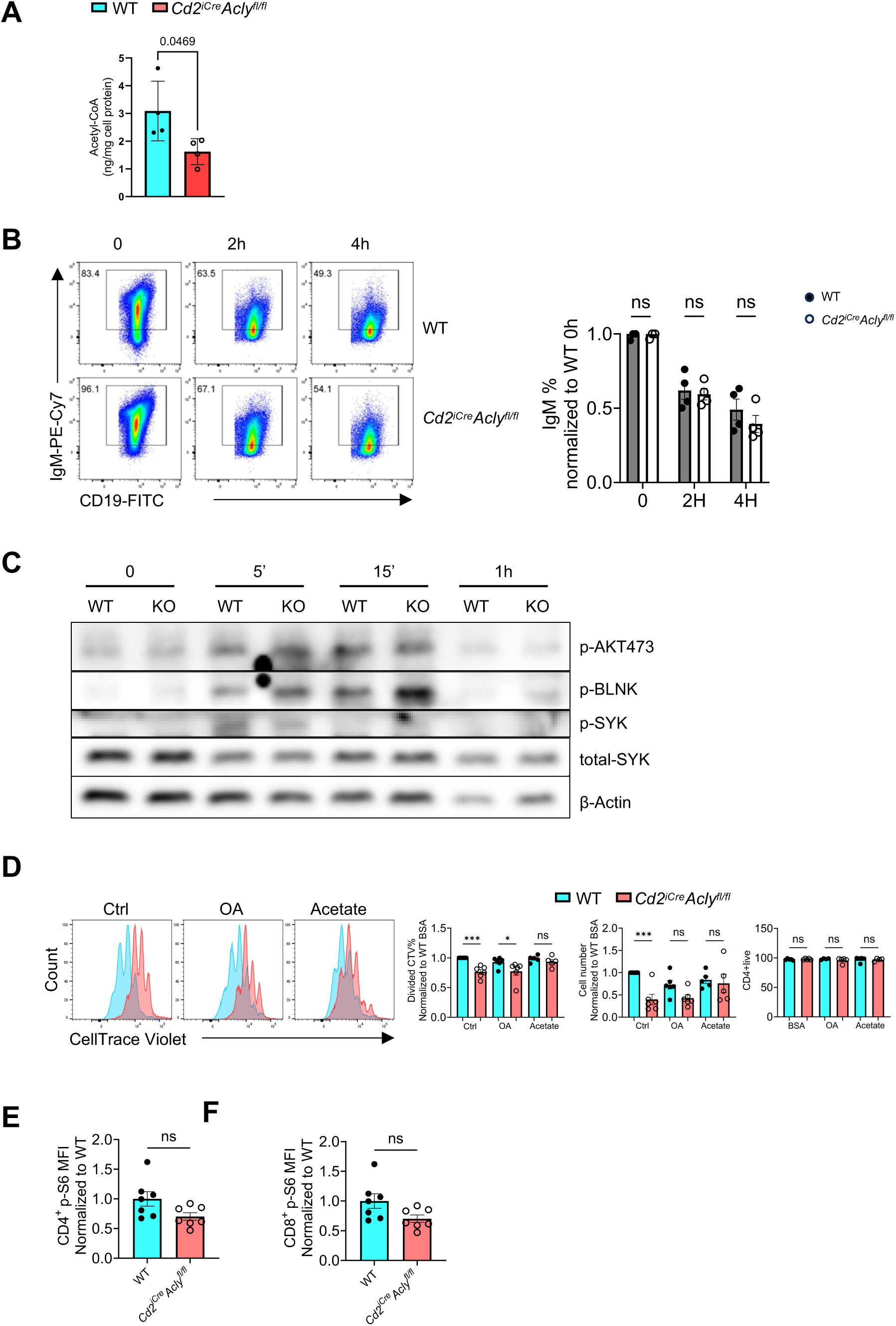
(A) Levels of total acetyl-CoA in B cells from WT and Cd2^iCre^Acly^fl/fl^ mice stimulated with LPS/IL4/BAFF for 16 h. (B) Left, B cell from WT and *Cd2^iCre^Acly*^fl/fl^ mice treated with 0.1 µg/mL anti-IgM for 0, 2 h and 4 h were labeled with IgM and CD19. Right, normalized IgM+ percentage. (C) B cells from WT and *Cd2^iCre^Acly*^fl/fl^ mice treated with 10 µg/mL anti-IgM at different time points. Immunoblot analysis of p-AKT473, p-BLINK, p-SYK and total SYK were performed on B cells, with β-Actin serving as the control. (D) Left, CD4^+^ T cells from spleens of WT and *Cd2^iCre^Acly*^fl/fl^ mice were labeled with CTV after treated with BSA (50 µM), OA (50 µM), or sodium acetate (5 mM) for 3 days. Right, normalized divided CTV and cell number of CD4+ T cells. (E-F) Normalized p-S6 MFI of CD4^+^ (F) and CD8^+^ (G) T cells after stimulated with anti-CD3/anti-CD28 for 4 h. *p* values were calculated using Student’s t test. ns, not significant, *p < 0.05, **p < 0.01, ***p < 0.001. Data are representative of at least three independent experiments (A, C). Error bars represent SEM.

**Supplementary Figure 3.**
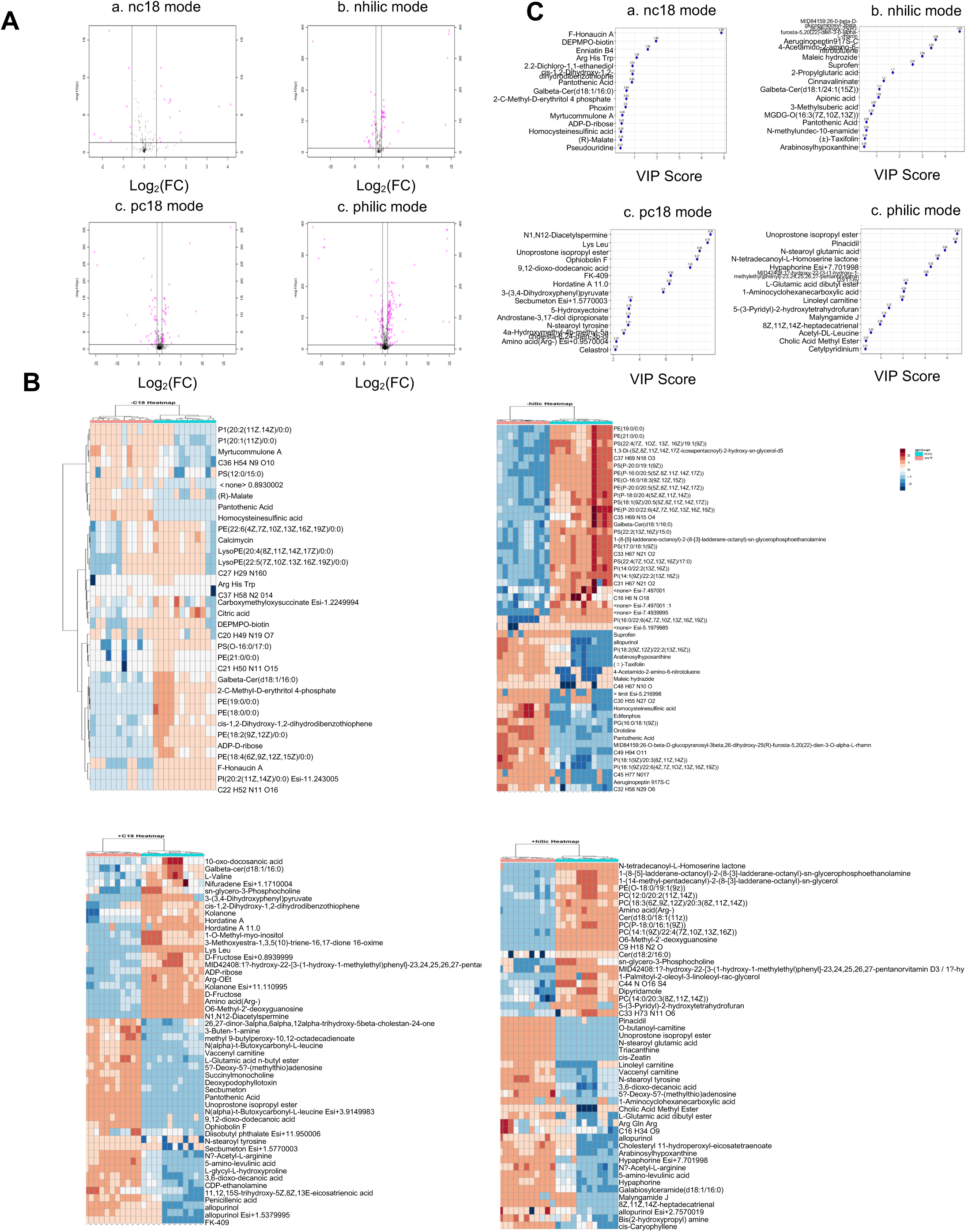
Untargeted metabolomic profiling of stimulated WT and KO B cells. (A) Volcano plots showing differentially abundant metabolites across the four acquisition modes: c18 chromatography in positive and negative ion modes (pC18 and nC18) and HILIC chromatography in positive and negative ion modes (philic and nhilic). (B) Heatmaps showing abundance patterns of differentially abundant metabolites across the four acquisition modes. (C) VIP score plots showing the top discriminative metabolites across the four acquisition modes.

**Supplementary Figure 4.**
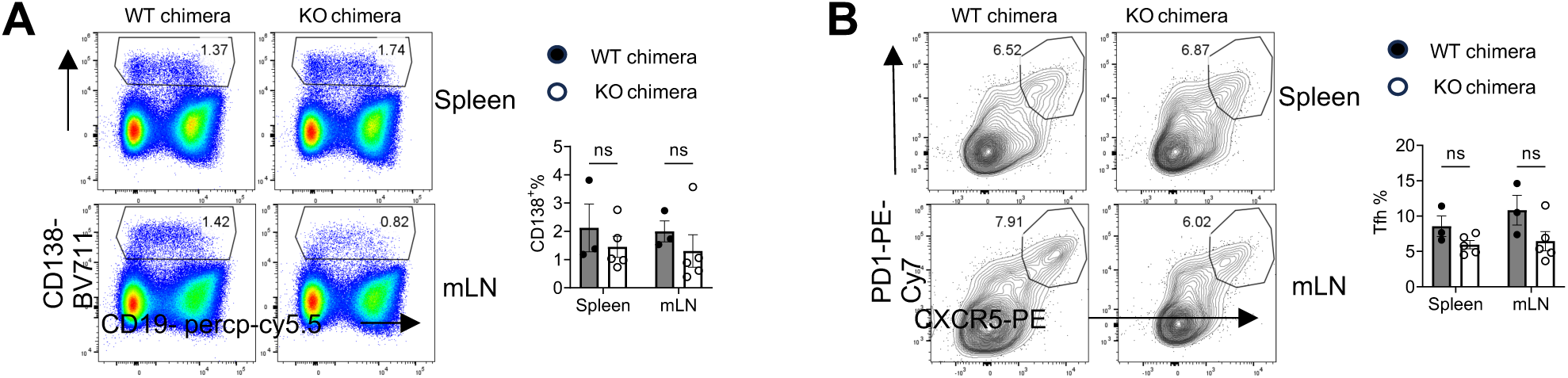
(A-B) μMT:WT chimera and μMT:*Cd2*^iCre^*Acly*^fl/fl^ chimera mice were infected with H1N1 PR8 influenza strain intranasally and were analyzed 14 days after infection. Left, CD19 and CD138 expression on lymphocytes from spleen or mediastinal lymph nodes (mLN) (A), and expression of PD-1 and CXCR5 among CD4^+^ T cell from spleen or mLN (B). Right, the frequency of CD138^+^ plasmablasts from spleen and mLN (A), and PD-1^+^CXCR5^hi^ Tfh cells from spleen and mLN (B). *p* values were calculated using student t-test or two-way ANOVA. ns, not significant, *p < 0.05. Error bars represent SEM.

**Supplementary Figure 5.**
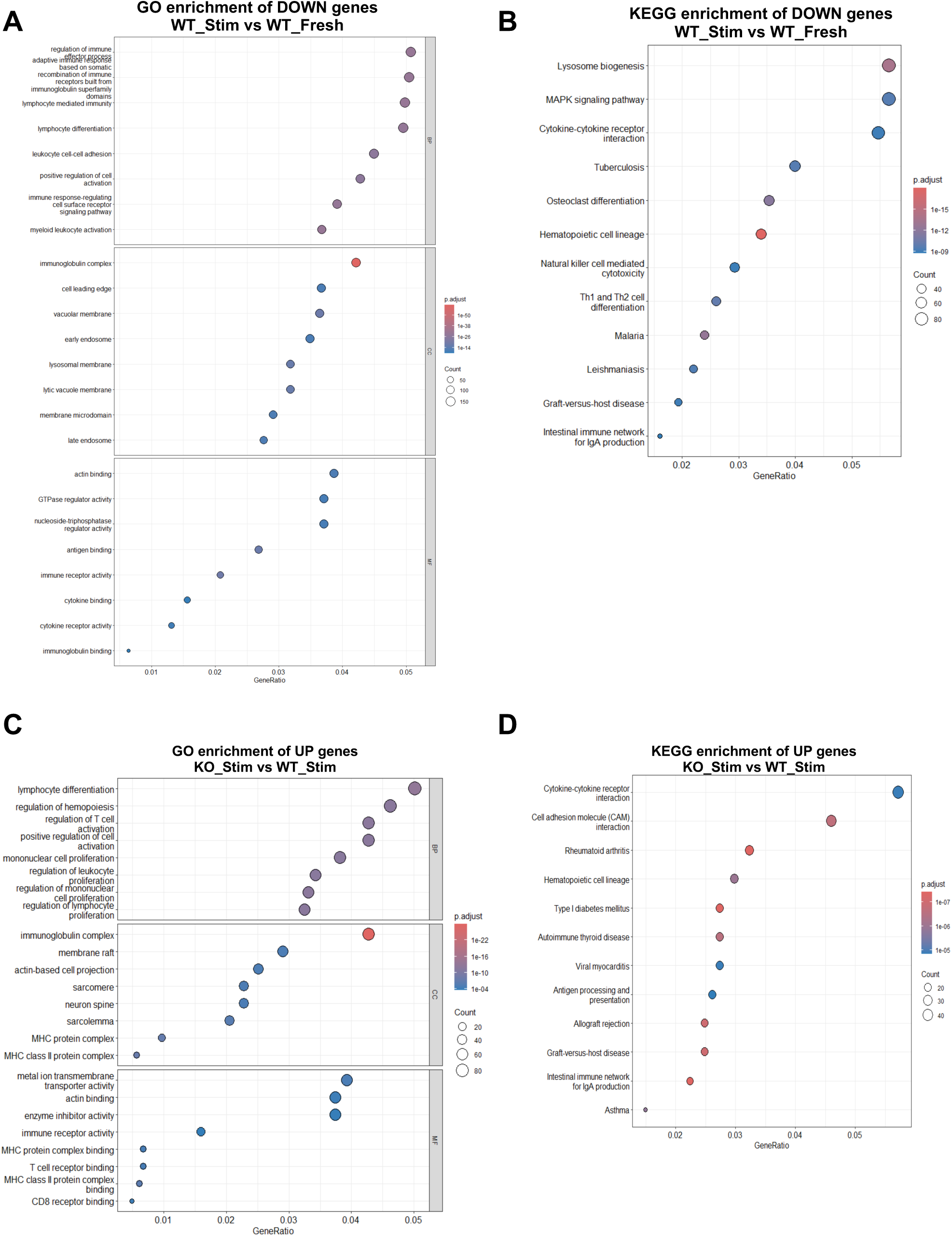
Functional enrichment analyses of differentially expressed genes. (A) Gene Ontology (GO) enrichment analysis of downregulated genes in WT_Stim versus WT_Fresh, including biological process (BP), cellular component (CC) and molecular function (MF) categories. (B) KEGG pathway enrichment analysis of downregulated genes in WT_Stim versus WT_Fresh. (C) GO enrichment analysis of upregulated genes in KO_Stim versus WT_Stim, including BP, CC, MF categories. (D) KEGG pathway enrichment analysis of upregulated genes in KO_Stim versus WT Stim.

**Supplementary Figure 6.**
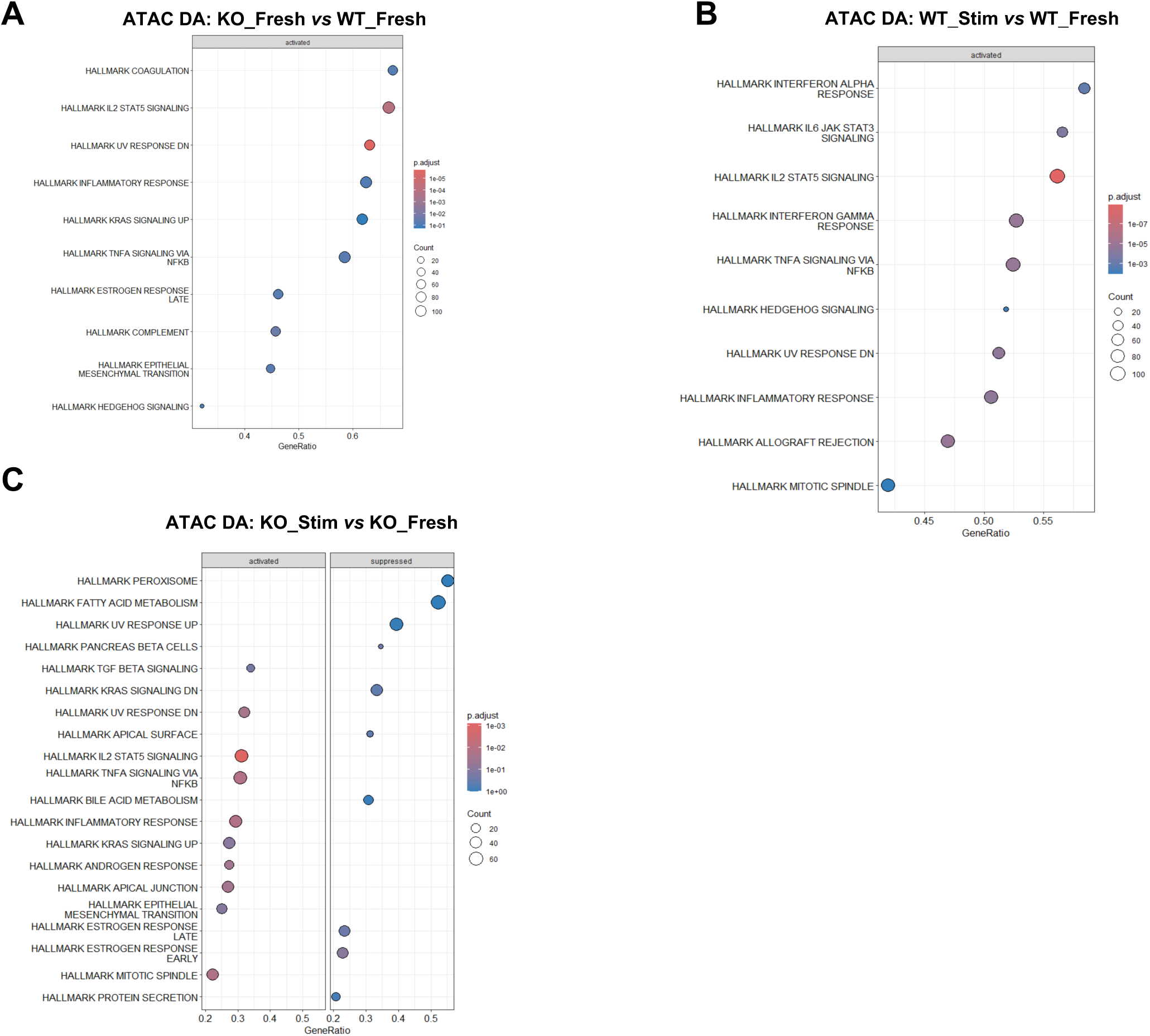
Functional and pathway enrichment analyses of differential chromatin accessibility. (A) GSEA of differential chromatin accessibility (DA) in KO_Fresh versus WT_Fresh. (B) GSEA of DA in WT_Stim versus WT_Fresh. (C) GSEA of DA in KO_Stim versus KO_Fresh.

